# Examining the effects of an anti-Salmonella bacteriophage preparation, BAFASAL, on *ex vivo* human gut microbiome composition and function using a multi-omics approach

**DOI:** 10.1101/2021.07.04.451072

**Authors:** Janice Mayne, Xu Zhang, James Butcher, Krystal Walker, Zhibin Ning, Ewelina Wójcik, Jarosław Dastych, Alain Stintzi, Daniel Figeys

**Author notes:** **Corresponding Authors**: D.F., and J.D.

## Abstract

*Salmonella* infections (salmonellosis) pose serious health risks to humans, usually via contamination in our food chain. This foodborne pathogen causes major food losses and human illnesses that result in significant economic impacts. Pathogens such as *Salmonella* have traditionally been kept at bay through the use of antibiotics, but antibiotic overuse within the food industry has led to the development of numerous multidrug-resistant bacterial strains. Thus, governments are now restricting antibiotic use, forcing the industry to search for alternatives to secure safe food chains. Bacteriophages, viruses that infect and kill bacteria, are currently being investigated and used as replacement treatments and prophylactics due to their specificity and efficacy. They are generally regarded as safe alternatives to antibiotics as they are natural components of the ecosystem. One example is BAFASEL, a commercial bacteriophage mixture that specifically targets *Salmonella* and is currently approved for use in poultry farming. However, when specifically used in the industry they can also make their way into humans through our food chain or exposure as is the case for antibiotics. In particular, agricultural workers could be repeatedly exposed to bacteriophages supplemented in animal feeds. To the best of our knowledge, no studies have investigated the effects of such exposure to bacteriophages on the human gut microbiome. In this study, we used a novel *in vitro* assay called RapidAIM to investigate BAFASAL’s potential impact on five individual human gut microbiomes. Multi-omics analyses, including 16S rRNA gene sequencing and metaproteomic, revealed that *ex vivo* human gut microbiota composition and function were unaffected by BAFASAL treatment providing an additional measure for its safety. Due to the critical role of the gut microbiome in human health and the known role of bacteriophages in regulation of microbiome composition and function, we suggest assaying the impact of bacteriophage-cocktails on the human gut microbiome as a part of their safety assessment.

**Graphical Abstract:** 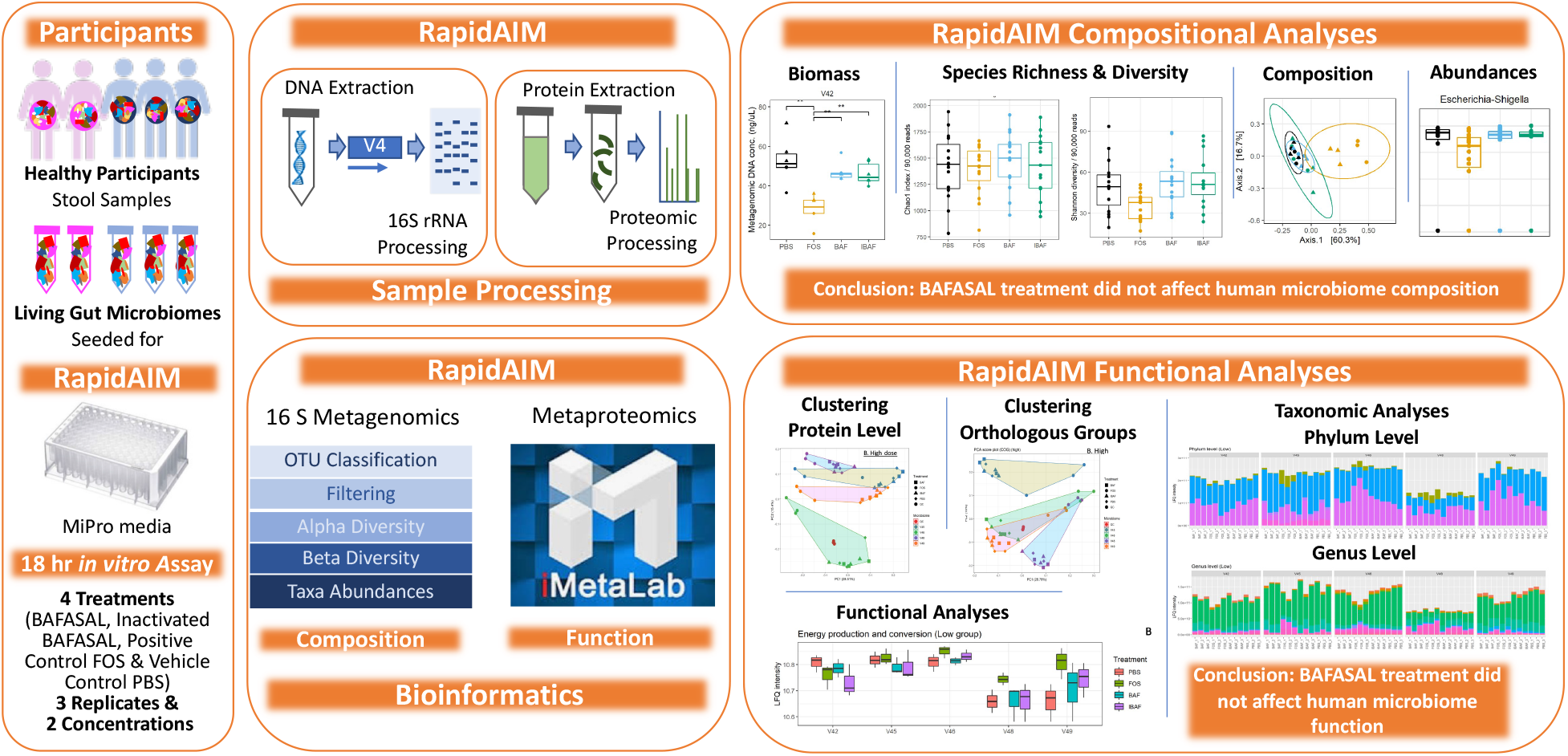

## 1. Introduction

As the world population increases, so does food demand. Plant and animal diseases can seriously impact food supplies and food safety resulting in food shortages and causing significant economic impacts. Zoonotic diseases present clear public health risks; in the USA, tens of thousands get sick each year with six out of 10 infectious diseases having a zoonotic origin [1]. Striking, the World Health Organization (WHO) estimates that consumption of contaminated food sickens 600 million people (or 10% of the world population), and results in 420,000 deaths [2].

Historically, most zoonotic diseases have been treated with antibiotics, with 73% of all antibiotics in the world now used in animal production [3]. The wide-spread use of antibiotics in the food industry was spurred on not only by increased food safety but by data showing increased meat production in animals receiving broad-spectrum antibiotics [4]. The appearance of multidrug resistant bacteria in our food chain is an unintended consequence from decades of over-using broad-spectrum antibiotics. In addition, humans are exposed to these excess antibiotics through several means; these antibiotics can be present in the animal products we consume, and can be simply discarded into the environment, with up to 90% of antibiotics excreted in their active forms from treated animals [5]. The rise of multidrug-resistant bacteria has become such a global concern that by 2022 the European Union (EU) will ban the prophylactic use of clinically relevant antibiotics in food production, with the exemption of veterinary prescriptions [6], and other countries are following suit. Without such interventions, antimicrobial resistance (AMR) will result in an estimated 10 million deaths by 2050 [5].

Nevertheless, the issue of food safety and the prevention of zoonotic diseases remain. Bacteriophage therapy is an alternative garnering increased interest by scientists and food agencies around the world. Discovered by Félix d’Hérelle and Frederick Twort in the early 1900s, the bacteriophages have been investigated for their therapeutic potential to treat pathogenic bacteria in humans, animals, and agriculture, especially in light of AMR [7]. Bacteriophages, or phages, are ubiquitous and abundant in nature, with numbers estimated above 10^31^ [8]. Indeed phages, outnumber their bacterial hosts by a factor of 10 times and regulate the bacterial populations they target by triggering bacterial lysis. Their ability to act directly on specific bacteria – via their recognition of unique receptor proteins - allows researchers to develop targeted therapies to fight against identifiable bacterial pathogens in our food chain and reduce the spread of foodborne illnesses in humans.

Bacteriophage therapy in the food industry has proven successful: Phages have been approved for use in food decontamination [9], as dietary supplements [10], and as environmental prophylaxis [11]. Importantly, no adverse effects were noted in trials as phages did not interact with eukaryotic cells and targeted only bacteria [11]. One of the most common foodborne illnesses, Salmonellosis, is caused by the bacterium *Salmonella. Salmonella* easily transfers from its primary hosts (commonly laying hens, pigs, turkeys, and broilers) to humans and is a common target for bacteriophage cocktails [12]. *Salmonella* is a genus of the family *Enterobacteriaceae* and contains two species: *S. enterica* and *S. bongori* [13]. Salmonellosis causes fever, sepsis, infection of tissues, and inflammation of the gastrointestinal tract. In America alone, *Salmonella* genera cause 1.35 × 10^6^ infections, resulting in 26,000 hospitalizations and 420 deaths each year [1]. In the EU, salmonellosis is the most common foodborne disease and *S.enterica* the most frequently reported pathogen. In 2020, a study profiled the effectiveness of an anti-*Salmonella* bacteriophage cocktail, BASFASAL, to prevent contamination of dried and liquid poultry feed *in vitro* and to reduce number of Salmonella in intestines of birds challenged with *Salmonella in vivo* [12].

The human gut microbiome consisting of bacteria, viruses, and fungi plays a critical role in health, as evidenced by perturbations in the microbiome composition and/or function that associated with a variety of diseases, including inflammatory bowel diseases (IBD) [14], cardiovascular disease [15], Parkinson’s and Alzheimer’s diseases [16, 17], anxiety and depression [18]. Bacteriophages are a normal part of the human microbiota and also outnumber bacteria in our gut by at least ten-fold. Most are temperate phages, and so induction of these prophages under various conditions could disturb microbiota balance [19]. Although bacteriophages are generally recognized as safe (GRAS) when used for pathogen control in food processing, including ready to eat foods and poultry, bacteriophages could interact with niche gut microbiota [19]. Effects could range from no effect to direct lysis and dysbiosis of commensal bacterial to indirect interactions of bacteriophage with commensal bacteria or prophages triggering lysis [19].

We have developed an *in vitro* microbiome culturing system (MiPro) coupled to a downstream bioinformatics toolbox collectively called RapidAIM (or rapid assay of an individual microbiome) that measures compositional and functional changes in the gut microbiome in response to dietary and therapeutic interventions [20, 21]. RapidAIM can measure the impact of products on the microbiota and can be used to stratify people in clinical studies. We have previously used this assay to show the effects of antibiotics and other drugs on the microbiome [20]. In this study, we used RapidAIM to investigate further the effects of BAFASAL; specifically, does BAFASAL phage introduction to the human microbiome affect its composition and function. We treated five individual adult-derived microbiomes with active and inactive mixtures of BAFASAL. We used both 16S rRNA gene sequencing and metaproteomic analyses to study changes in bacterial composition and function. This study, the first of its kind, has important implications for how we measure food safety and maintain the safety of workers in the food processing industry exposed to these preparations.

## 2. Materials and Methods

### 2.1. BAFASAL^®^ production

Liquid BAFASAL preparation was produced according to Wójcik *et al*. [12]. Briefly, bacteriophages included in BAFASAL (BAF) were amplified separately in culture, reaching a titer of about 1 × 10^9^ PFU/mL. Phage-containing culture fluid was separated from bacterial debris by filtration, and final titers assessed using double agar overlay plaque assay. The final preparation was completed with sterile water, resulting in a 1 × 10^8^ PFU/mL titer. Inactive BAFASAL (IBAF) was prepared by heating this preparation at 120°C for 30 min.

### 2.2. Stool sample preparation

The research ethics board protocol (# 20160585-01H) for human stool sample collection was approved by the Ottawa Health Science Network Research Ethics Board at the Ottawa Hospital. Stool samples were obtained from five healthy volunteers (age range 27-36 years; three men and two women). Exclusion criteria for participation were irritable bowel syndrome, inflammatory bowel disease, or diabetes diagnosis; antibiotic use or gastroenteritis episode in three months preceding collection; use of pro-/pre-biotic, laxative, or anti-diarrheal drugs in the last month preceding collection; or pregnancy. Participants collected stool into a 50 mL Falcon tube containing 15 mL of sterile phosphate-buffered saline (PBS; pH 7.6) and 10% (v/v) glycerol pre-reduced with 0.1% (w/v) L-cysteine hydrochloride. Samples were weighed, transferred into an anaerobic workstation (5% H_2_, 5% CO_2_, and 90% N_2_ at 37°C), homogenized to 20% (w/v) in the same pre-reduced buffer mixture and filtered using sterile gauzes to remove large particles to obtain the fecal inocula. Fecal inocula, as proxies for gut microbiomes, were stored at −80°C until used in RapidAIM.

### 2.3. Culturing of Microbiota and treatments

Fecal inocula were thawed at 37°C and inoculated at a concentration of 2% (w/v) into a 96-well deep-well plate containing 1 mL MiPro optimized sterile and pre-reduced culture medium (2.0 g L^−1^ peptone water, 2.0 gL^-1^ yeast extract, 0.5 g L^-1^L-cysteine hydrochloride, 2 mL L^-1^ Tween 80, 5 mg L^-1^ hemin, 10 μL L^-1^ vitamin K1, 1.0 g L^-1^NaCl, 0.4 g L^-1^ K_2_HPO_4_, 0.4 g L^-1^ KH_2_PO_4_, 0.1 g L^-1^ MgSO_4_·7H_2_O, 0.1 g L^-1^ CaCl_2_·2H_2_O, 4.0 g L^-1^NaHCO_3_, 4.0 g L^-1^ porcine gastric mucin, 0.25 g L^-1^ sodium cholate, and 0.25 g L^-1^ sodium chenodeoxycholate) as per Li *et al*. in an anaerobic workstation [20, 21]. BAF and IBAF were added at 2% (v/v) and 5% (v/v) to the media. For BAF, this equated to 2 × 10^6^ and 5 × 10^6^ PFU/mL, respectively. Fructo-oligosaccharide (FOS) was added at 2% (v/v) and 5% (v/v) from a 250 mg/mL stock as a positive control and PBS (1X; pH 7.6) added as vehicle control at 2% (v/v) and 5% (v/v). Following the addition of inoculants and compounds, the plates were covered with vented sterile silicone mats and shaken at 500 rpm with a digital shaker (MS3, IKA, Germany) at 37°C for 18h in the anaerobic chamber. Treatments were randomized on the 96-well plates for each participant.

### 2.4. Sample processing

Sample processing was done as per our reported sample processing workflows [20-22]. Briefly, following 18 hrs of culturing, samples were transferred from 96-well plates to individual 1.5 mL Eppendorf tubes. Sample order were randomized for processing. Samples were transferred out of the anaerobic station onto ice and centrifuged at 14000*g* at 4°C for 20 min to pellet the microbiota. Supernatants were removed, and the pellets were resuspended in 1 mL cold 1X PBS (pH7.6). They were subsequently centrifuged at 300*g* at 4°C for 5 min to remove debris. With samples on ice, supernatants were transferred into new 1.5mL Eppendorf tubes for two additional rounds of debris removal as above. Supernatants were transferred to 1.5mL Eppendorf tubes and centrifuged at 14000*g* at 4°C for 20 min to pellet the microbiota. Supernatants were removed, and the pelleted bacterial cells were resuspended and washed two times with 1 mL cold 1X PBS (pH7.6), pelleting the cells after each wash at 14000*g* for 20 min at 4°C. Before the final spin, resuspended bacterial samples were equally divided into new 1.5mL Eppendorf tubes. Harvested bacterial cell pellets were stored at −80°C prior to protein extraction or DNA extraction.

### 2.5. Metaproteomic sample processing and LS-MS/MS analyses

Microbial pellets were resuspended in 100 μL of 8 M urea and 4% (w/v) sodium dodecyl sulfate (SDS) in 100 mM Tris-HCl (pH 8.0) plus Roche cOmplete™ Mini tablets. Lysis was completed with sonication (Q700 Qsonica, USA) at 8°C, 50% amplitude (15.6 watts/sample), and sixty cycles of 10 s ultrasonication and 10 s cooling down. Samples were centrifuged at 16000*g* for 10 min at 9°C. Supernatants were transferred to new 1.5 mL Eppendorf tubes and proteins in the supernatant were precipitated with 5X volume of ice-cold protein precipitation buffer (acidified acetone/ethanol) overnight at −20°C. Proteins were pelleted by centrifugation at 16000*g* for 20 min at 4°C and washed 3X in ice-cold acetone. Proteins were resuspended in 100 μL of 6M urea in 50 mM ammonium bicarbonate (ABC; pH 8.0) with sonication as above. Protein concentrations were measured in triplicate using Bio-Rad’s detergent compatible, DC Protein Assay (USA). Trypsin digestion was carried out as described in Zhang *et al*. [23]. For each sample, 50 μg of protein was reduced with 10 mM dithiothreitol (DTT) at 56°C for 30 min and alkylated in the dark with 20 mM iodoacetamide (IAA) at room temperature for 45 min, followed by 10X dilution into 50 mM ABC (pH 8.0) and tryptic digestion at 37°C for 20 hr with shaking using 1 μg of trypsin (Worthington Biochemical Corp., Lakewood, NJ) per 50 μg protein. Trypsin digestion was stopped by addition of 10% (v/v) formic acid (FA) to pH 2-3. Acidified, digested peptides were desalted using 10 μm C18 column beads according to our laboratory standard protocol[20], dried via vacuum centrifugation, and resuspended in 100 μL 0.1% (v/v) FA for tandem mass spectrometry (MS/MS) analyses. As quality control (QC) samples, to be assessed repeatedly throughout our MS/MS runs, 10 peptide samples were selected at random and combined. Two microlitres (1 μg peptides) were loaded in a randomized order for liquid chromatography tandem mass spectrometry (LC-MS/MS) onto an Agilent 1100 Capillary LC system (Agilent Technologies, San Jose, CA) and a Q Exactive mass spectrometer (ThermoFisher Scientific Inc.). Peptides were separated on a tip column (75 μm inner diameter × 50 cm) packed with reverse phase beads (1.9 μm/120 Å ReproSil-Pur C18 resin, Dr. Maisch GmbH, Ammerbuch, Germany) using a 90-min gradient from 5 to 30% (v/v) acetonitrile at a 200 nL/min flow rate. 0.1% (v/v) FA in water was used as solvent A, and 0.1% FA in 80% acetonitrile was used as solvent B. The MS scan was performed from 300 to 1800 m/z, followed by data-dependent MS/MS scans of the 12 most intense ions, a dynamic exclusion repeat count of two, and repeat exclusion duration of 30 s were used. The resolutions for MS and MS/MS were 70,000 and 17,500, respectively.

### 2.6. Metaproteomic data analysis

Mass spectrometry proteomics data were deposited to the ProteomeXchange Consortium via the PRIDE partner repository. Peptide/protein identification and quantification, taxonomic assignment, and functional annotations were done using MetaLab (version 1.1.0) [24]. Briefly, MetaLab automates an iterative database search strategy called MetaPro-IQ, described in Zhang *et al*. (2016). The MetaPro-IQ search was based on a human gut microbial gene catalog containing 9,878,647 sequences from http://meta.genomics.cn/. In MetaLab, a spectral clustering strategy [24] was used for database construction from all raw files. Then the peptide and protein lists were generated by applying strict filtering based on a false discovery rate (FDR) of 0.01, and quantitative information for proteins was obtained with the maxLFQ (label-free quantification) algorithm on MaxQuant (version 1.5.3.30). Carbamidomethyl (C) was set as a fixed modification, and oxidation (M) and N-terminal acetylation (Protein N-term) were set as variable modifications. The matching between runs option was used. Instrument resolution was set as “High-High”. Quantified protein groups were filtered according to the criteria that the protein appeared in >50% of the microbiomes for each treatment. Protein group LFQ intensities were then log_2_-transformed. Functional annotations of protein groups, including COG (Clusters of Orthologous Groups) information, were obtained using MetaLab. Functional responses, including hierarchical clustering, heatmaps, and principal component analyses (PCA) were analyzed and visualized using R (version 4.0.2) or Shiny apps on imetalab.ca.

### 2.7. Metagenomic DNA Extraction and 16S rDNA-V4 amplicon sequencing

Metagenomic DNA extraction from samples (RapidAIM cultures and stools) and V4-16S rRNA gene library construction and sequencing were done as previously described [25, 26]. Samples were extracted in the same randomized order as in the metaproteomic analysis. Briefly, metagenomic DNA was extracted using beads beating, extracted DNA normalized, and the V4-16S rRNA gene PCR amplified. Positive controls included a ZymoBiotics community standard (Cat# D6300) that was processed alongside the study samples. Negative controls included extraction blanks and DNA-free water (Ambion Cat#: AM9935) as the template for the V4-16S rRNA PCR. We processed BAF and IBAF preparations through our extraction/sequencing pipeline and also used pure BAF/IBAF as a template to our V4 amplicon PCR to identify potential contaminating bacteria in the BAF/IBAF preparations. V4 PCR amplicons were normalized by mass, pooled together, sized selected, and quantified using an Agilent Bioanalyzer. The pooled library was subsequently templated and sequenced using an IonChef and Ion Torrent Proton sequencer using the manufacturer’s recommended protocol. The samples were partitioned between the two sequencing chips in a randomized fashion. Two separate V4-16S rRNA amplicon libraries were constructed for the input stool samples and included on each sequencing chip to assess reproducibility.

### 2.8. Processing for 16S rRNA sequencing and analyses

Raw sequencing reads were quality filtered and demultiplexed prior to OTU (operational taxonomic unit) picking as previously described. The demultiplexed reads are available at the NCBI Sequence Read Archive (http://www.ncbi.nlm.nih.gov/sra). QIIME 1.9.1 was used to identify OTUs using a closed reference strategy against the SILVA 119 database, and the resulting data analyzed in R. Potential contaminants were removed with the decontm package using V4 amplicon concentrations prior to pooling. OTUs were subsequently filtered to only keep those with ≥2 counts at least 5% of the final sample dataset, and samples were rarefied to 90,000 reads prior to analysis. Samples with less than 90,000 reads were discarded. Differences in alpha diversities (Chao1 index and Shannon diversity) were assessed using a Kruskal-Wallis test with Dunn’s posthoc test. Beta diversities were assessed using the Bray-Curtis dissimilarity, weighted Unifrac distance, and unweighted Unifrac distance. The impact of BAF or IBAF on beta diversity clustering was assessed with adonis function from vegan by pooling the samples for each stool donor. Potentially differentially abundant taxa between the PBS and BAF/IBAF cultures were assessed using MaAsLin2 controlling for stool donors using the default parameters. Results were considered significant with a *p-value* <0.05 and, where appropriate, after controlling for multiple hypothesis testing using the Benjamini and Hochberg approach.

## 3. Results

### 3.1. Experimental Set-Up

In this study, we used our well-established and optimized RapidAIM (<u>Rapid</u> <u>a</u>ssay of an <u>i</u>ndividual <u>m</u>icrobiome) workflow to carry out both metaproteomic and metagenomic analyses of five microbiomes following *in vitro* treatment with the BAFASAL commercial bacteriophage mixture (Figure 1). Microbiotas were harvested from the feces of five participants (3 men and 2 women) as proxies of their gut microbiota. In brief, microbiota samples were derived from homogenized human feces and stabilized in 20% (w/v) anaerobic, pre-reduced 1X PBS/10% (v/v) glycerol/0.1% (w/v) L-cysteine. These samples were inoculated to 2% (w/v) in 1 mL of Mi-Pro optimized culture media and incubated under anaerobic conditions at 37°C for 18 hrs. We have shown that this *in vitro* assay maintains both microbiome composition and function over 24 hrs in culture [21]. To evaluate the potential effects of BAFASAL on human gut microbiota, we compared (1) BAFASAL-treated microbiota (BAF) to those microbiotas exposed to (2) heat-inactivated BAFASAL (IBAF), (3) positive control fructo-oligosaccharide (FOS), and (4) vehicle phosphate-buffered saline (PBS; pH 7.6). Samples were cultured in triplicate for each condition and at 2% (v/v) and 5% (v/v) concentrations as described in materials and methods. Following 18 hr culturing, the bacterial cells were pelleted, DNA extracted for metagenomic analyses using 16S rRNA gene-based sequencing, and proteins extracted and digested for metaproteomic analysis, as described in materials and methods. We used 16S rRNA gene amplicon data to analyze compositional changes and metaproteomics to focus on functional changes following microbiome treatments.

**Figure 1.**
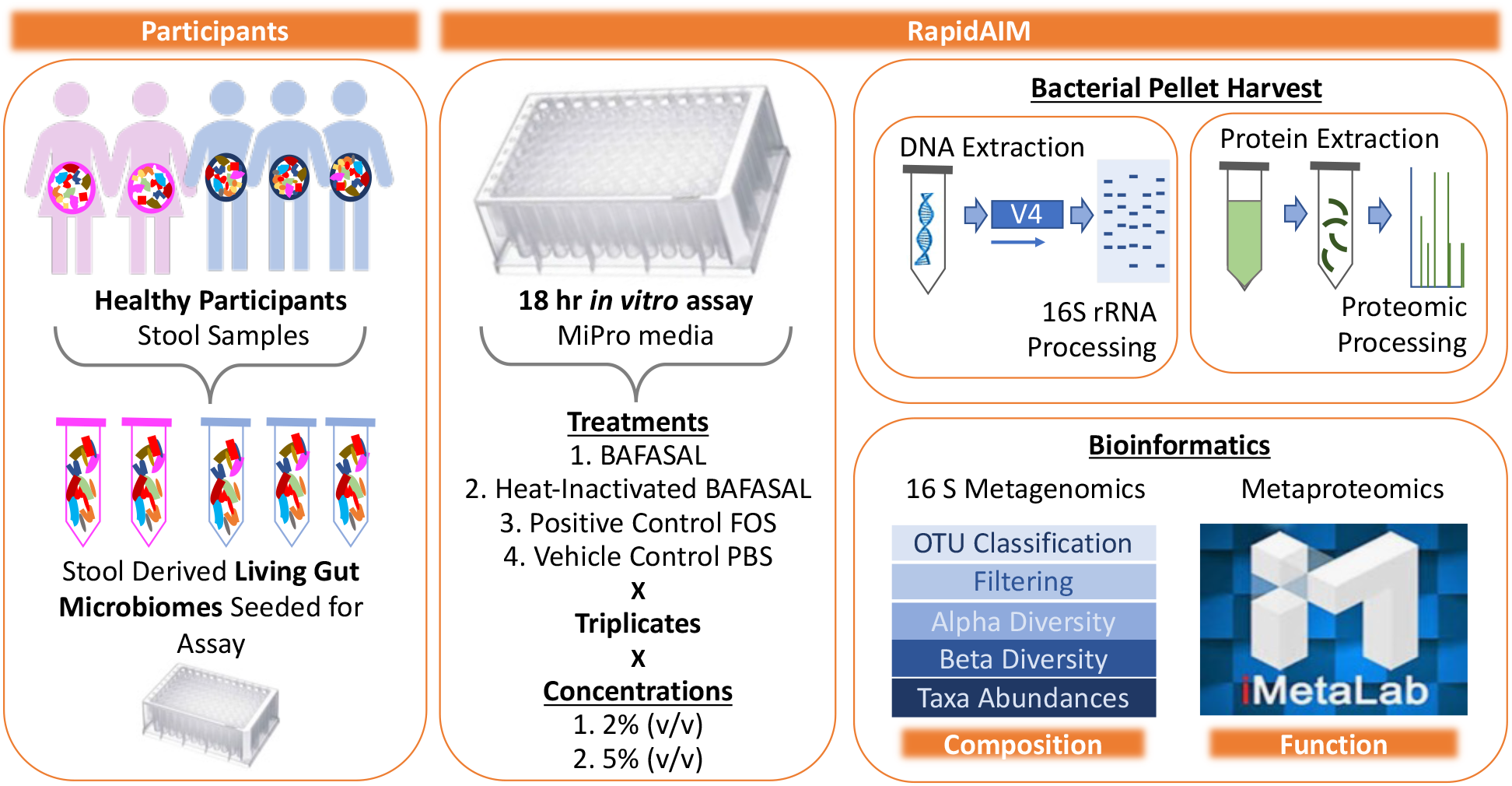
Experimental Set-Up. Microbiotas were harvested from the feces of five participants (3 men and 2 women) as proxies of their gut microbiota. Samples were derived from homogenized human feces stabilized in 20% (w/v) anaerobic, pre-reduced 1X PBS/10% (v/v) glycerol/0.1% (w/v) L-cysteine. Samples were inoculated to 2% (w/v) in 1 mL of Mi-Pro optimized culture media and incubated under anaerobic conditions at 37°C for 18 hrs. To evaluate the potential effects of BAFASAL on human gut microbiota, we compared (1) BAFASAL-treated microbiota (BAF) to those microbiotas exposed to (2) heat-inactivated BAFASAL (IBAF), (3) fructo-oligosaccharides as a positive control (FOS), and (4) phosphate-buffered saline (PBS; pH 7.6) as a negative control in triplicate and at two concentrations, as defined in materials and methods. Following culturing, microbial cells were pelleted and frozen until further analysis. DNA was extracted for metagenomic analyses using 16S rRNA gene-based sequencing and proteins extracted and digested for metaproteomic analysis, as described in materials and methods. Bioinformatic analysis was used to determine microbiome compositional and functional changes in response to treatment.

### 3.2. Microbial biomass

We firstly assessed whether BAFASAL addition would impact the ability of human microbiotas to grow *in vitro*. As a gross measure for tracking increases in microbial biomass, we measured the amount of metagenomic DNA and protein extracted for each individual under the different treatments (Figure 2). For all individuals, biomass did not differ following either BAF or IBAF treatments. Nor did these treatments differ when compared to the PBS vehicle control for our positive treatment FOS. In contrast, almost all individuals showed reduced bacterial biomass upon FOS treatment. This biomass reduction is likely due to the production of short-chain fatty acids that acidified the medium and inhibited specific microbes’ growth.

**Figure 2:**
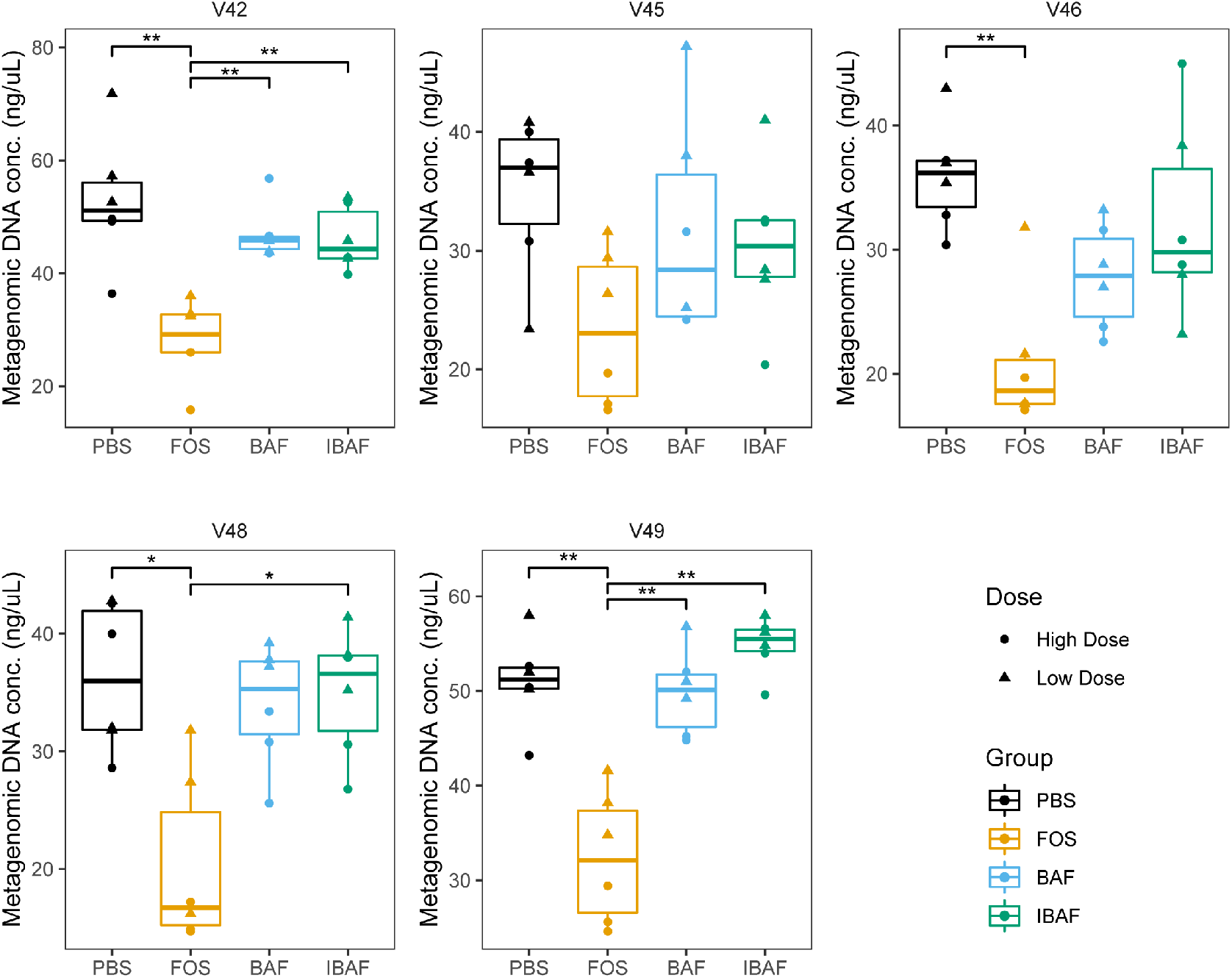
BAFASAL treatment does not impact microbial biomass after 18 hr of *in vitro* growth. Microbial biomass expressed as the quantity of metagenomic DNA (ng/uL) extracted for each donor and each treatment. Donors are plotted separately, and the high/low treatments combined. Statistical differences were assessed using Kruskal-Wallis test with Dunn’s multiple comparison post-hoc test. * p<0.05, ** p<0.01.

### 3.3. Compositional Analyses

#### 3.3.1. RapidAIM and sequencing library reproducibility

We assessed the reproducibility of the sequencing reactions by comparing the microbiome profiles of the stool samples that were sequenced on both chips for high and low doses. Principal coordinate analysis (PCoA) using the Bray-Curtis dissimilarity revealed that the individual stools clustered tightly together in a donor-dependent manner (Supplemental Figure 1A) with essentially no separation between the two sequencing chips. Moreover, we assessed the reproducibility of the RapidAIM culturing assay by comparing the PBS replicates between the high/low dose conditions as these would be expected to show few differences. This analysis revealed little separation between the two culturing conditions (Supplemental Figure 1B), again with the samples segregating primarily by an individual. Finally, our commercial standards revealed that we could detect all the genus present in this mix and could distinguish between the closely related *Escherichia coli* and *Salmonella enterica* present in the standard (Supplemental Figure 1C).

#### 3.3.2. Microbial diversity analyses

We next evaluated whether BAFASAL impacted microbial alpha diversity. Species richness (Figure 3A) and diversity (Figure 3B) did not differ significantly among BAF, IBAF, and PBS treatments at either dosing level. However, FOS treatment lowered the overall diversity at high concentrations but had no impact on overall species richness.

**Figure 3:**
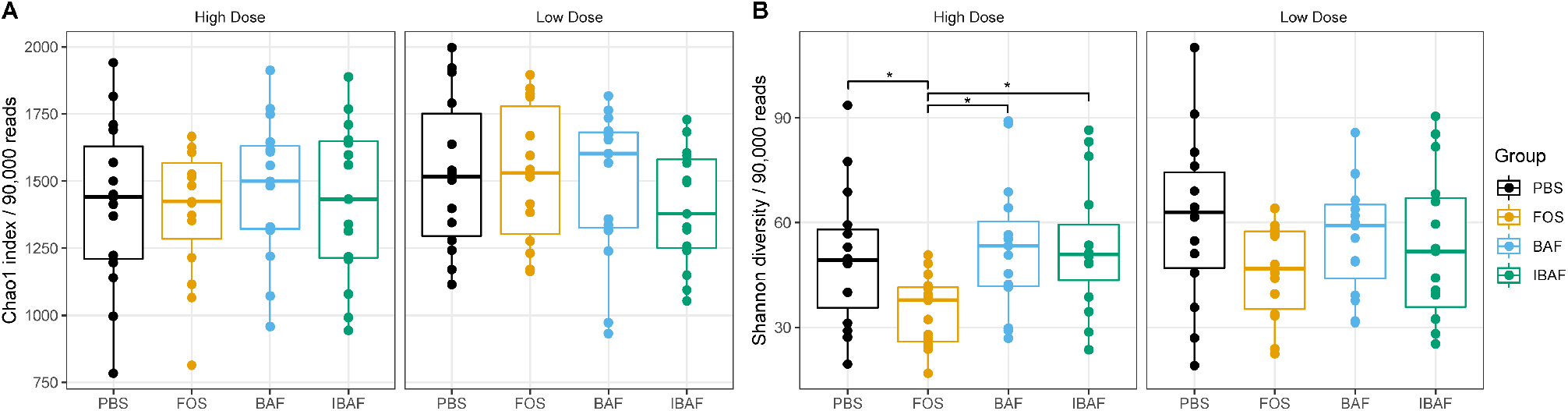
BAFASAL treatment does not impact microbial alpha diversity. Species richness (A) and diversity (B) were assessed for each treatment group. There was no difference in between the groups except for lowered species diversity in the high FOS concentration. Statistical differences assessed using a Kruskal-Wallis test with Dunn post-hoc test. * p<0.05, ** p<0.01.

We subsequently compared the microbiota beta diversity profile for all RapidAIM assayed samples (BAF, IBAF, PBS, and FOS, at both high and low doses; Figure 4). This analysis revealed that samples from a given individual formed their own cluster regardless of treatment, underscoring each individual’s highly personalized microbiota and the need to assess microbial responses on a personalized level.

**Figure 4:**
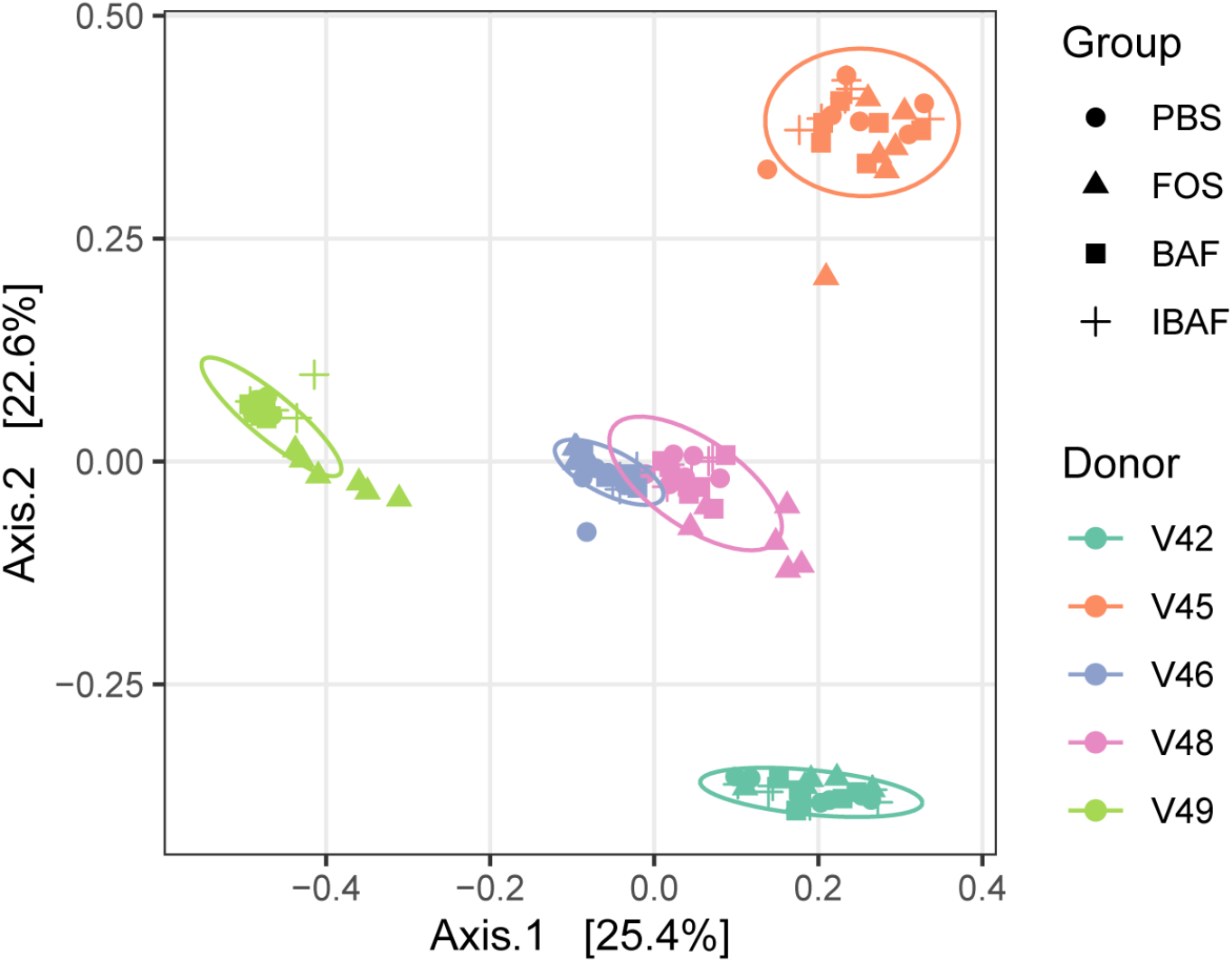
Principal coordinate analysis of the RapidAIM cultures reveal the highly personalized nature of human microbiomes. All RapidAIM assay microbiota results were analyzed using Principal coordinate analysis of the Bray-Curtis dissimilarity. This analysis revealed that each donor formed its own cluster, demonstrating the highly personalized nature of human microbiomes and the requirement for high-throughput in vitro assays to assess microbial responses on a personalized level.

Comparing the microbiota profiles for each individual’s treatments separately revealed that BAF and IBAF treatments did not result in major changes in the overall microbial community for any of the participants (Figure 5). This is in contrast with FOS treatment, which resulted in clear shifts in microbiota community composition for each individual. Notably, the lack of separation between the BAF, IBAF, and PBS communities persisted even after removing the FOS samples (Supplemental Figure 2) or when assessing the microbial communities using either weighted or unweighted Unifrac distances (data not shown).

**Figure 5:**
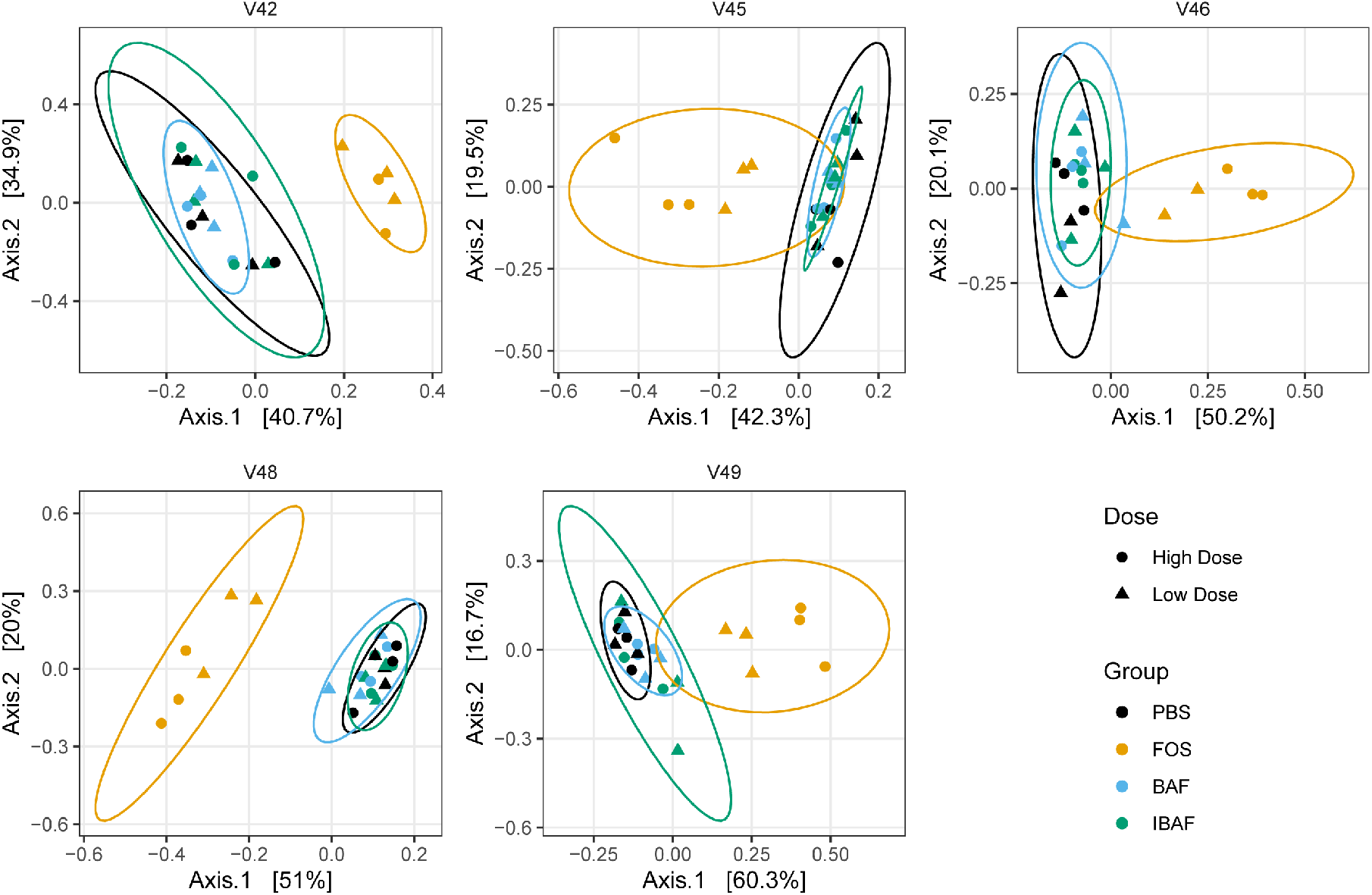
Principal coordinate analysis of RapidAIM cultures from each donor reveals that BAFASAL treatment has minimal impact on microbial composition. RapidAIM assays from each donor were analyzed separately using principal coordinate analysis of the Bray-Curtis dissimilarity and plotted as separate panels. This analysis reveals the large impact that FOS treatment has on patient microbiotas that could be obscured by high interpatient variability (Figure 4). There were no apparent differences between the PBS and BAFASAL/inactivated BAFASAL treatments.

#### 3.3.3 Compositional Differences

We could not identify significantly differentially abundant taxa between the BAF, IBAF, and PBS treated samples. This was true whether analyzing the high and low dosing regimens separately or by pooling the two dosing groups together. As BAF specifically targets *Salmonella*, we then focused on the abundances of related genera in the Enterobacteriaceae family. There were no apparent differences between the dominant Enterobacteriaceae genera in the BAF, IBAF, and PBS treated cultures (Supplemental Figure 3). In contrast with the above results, there were differences between the PBS and FOS treated samples that mirror previously reported results, such as increased *Bifidobacterium* genera (data not shown). Notably, and as expected, FOS treatment reduced the levels of several Enterobacteriaceae family members, such as *Escherichia* genera.

### 3.4. Functional analyses

Metaproteomic functional analyses were based on data from the MS/MS spectra collected with an average identification rate of 40.77% with 1,364,000 MS/MS submitted and 556,283 identified. In total, 65,760 peptides were identified and quantified that mapped to 16,064 proteins, across the 60 samples.

#### 3.4.1. Metaproteomic Assessed Responses to BAFASAL

To examine similarities and differences in protein expression following BAF, IBAF, PBS, and FOS treatments of the five microbiomes, we applied hierarchical cluster analyses for proteins identified in >50% of all samples (Q50) (Supplemental Figure 4, Panels A (low dose) and B (high dose)). As expected, quality control (QC) samples clustered together. While an individual’s microbiome samples clustered together, clustering did not occur among BAF, IBAF, and PBS treated microbiomes for an individual’s samples at either dose. Meanwhile, our positive control FOS induced significant shifts in protein abundances for each individual’s microbiome resulting in their clustering at both low and high doses. In fact, high-dose FOS treatment shifted protein expression patterns such that FOS treated microbiomes between individuals clustered together and moved further away from their corresponding control-treated PBS microbiomes (Panel B).

Principal component analysis (PCA) of those quantified protein groups (Q50) revealed that microbiome samples from an individual clustered together, showing again interindividual microbiome differences (Figure 6). No clustering occurred among BAF, IBAF and PBS treated microbiomes for any individual and at either low (Panel A) or high (Panel B) doses while QC samples clustered tightly. At low dose, FOS treated microbiomes formed a sub-cluster within an individual’s larger cluster (Panel A). At higher doses an individual’s sample treated with FOS separated from its BAF, IBAF and PBS treated samples along principal component axes 1, again indicating the variations induced by high dose FOS treatments were greater than intermicrobiome variations (Panel B).

**Figure 6.**
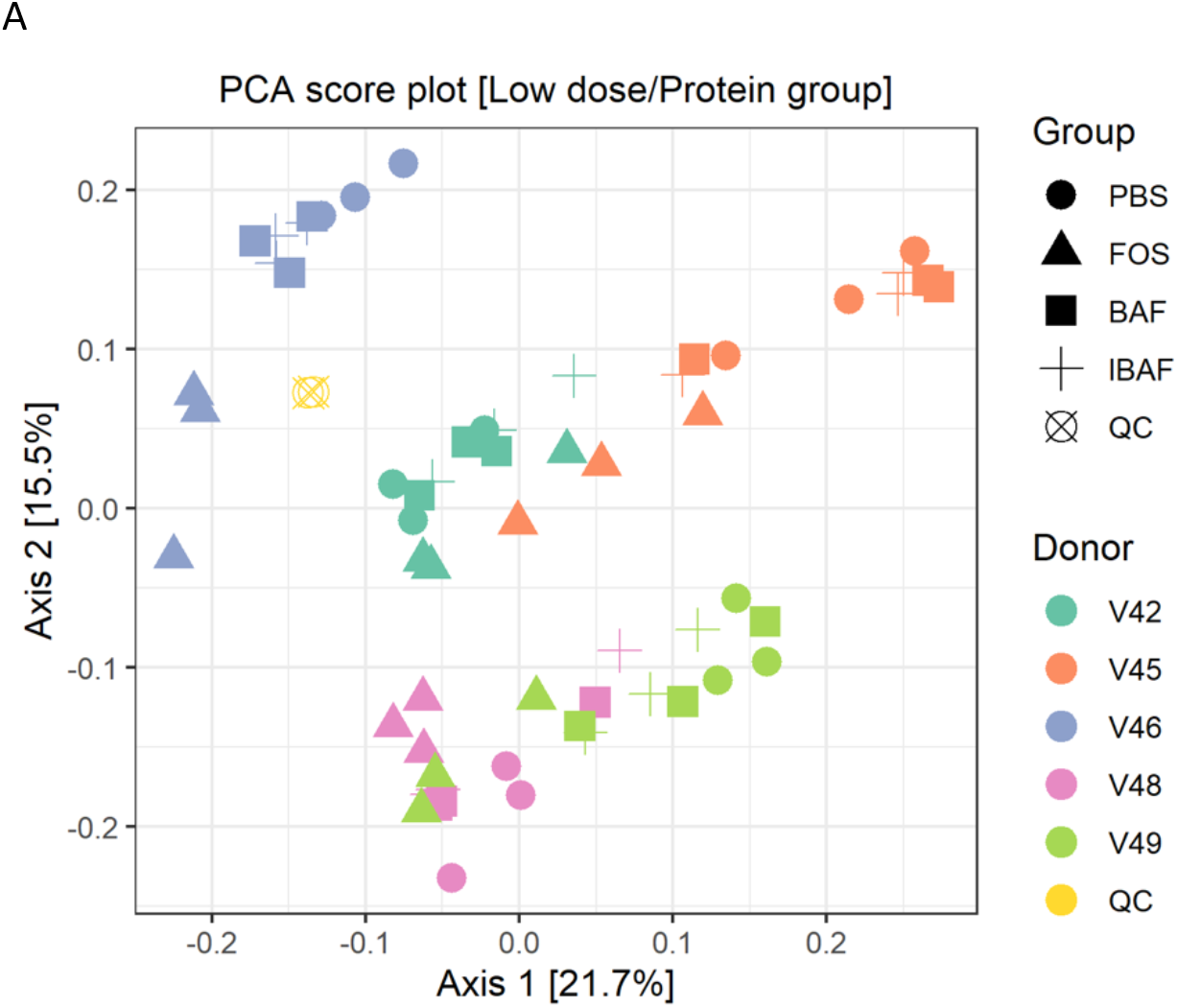
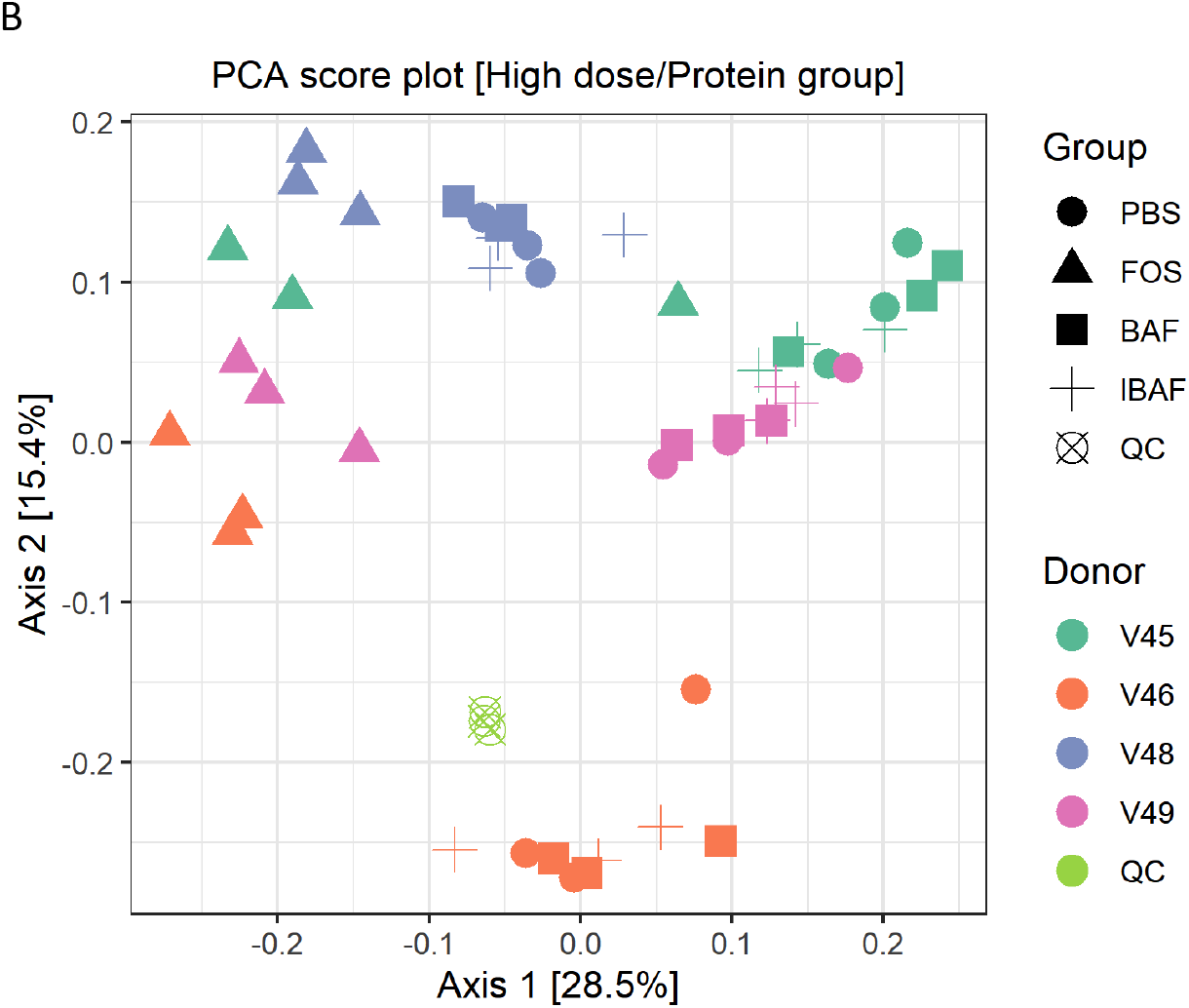
Principal component analyses of protein groups demonstrate the unique pattern of an individual’s protein expression profile and absence of BAFASAL impact. Panels A and B show principal component analyses (PCA) at the protein level per sample and for low and high doses, respectively. Colors indicate volunteer samples and shapes treatment as shown to legends on the right of each panel. Note: Only those protein groups that have non-zero values in >50% of samples were used for analyses (Q50).

#### 3.4.2 Functional Responses to BAFASAL

We studied the functional distribution of identified microbiome proteins and any changes induced with treatment via functional hierarchical clustering (Supplemental Figure 5) and PCA (Figure 7). Functional annotation of the protein groups using Clusters of Orthologous Groups (COGs) database did not reveal functional changes among BAF, IBAF, and PBS treated microbiomes for an individual and at either low (Panels A) or high dose (Panels B) treatment. Quality control samples clustered tightly together as expected. Similar to clustering observed for protein level analyses, at low doses, FOS-treated microbiomes formed a sub-cluster usually within an individual’s cluster (Figure 7, Panel A). Samples treated with FOS at higher doses separated from other samples along principal component axis 1, indicating again that the functional variations induced by high dose FOS were greater than intermicrobiome functional variations (Figure 7, Panel B).

**Figure 7.**
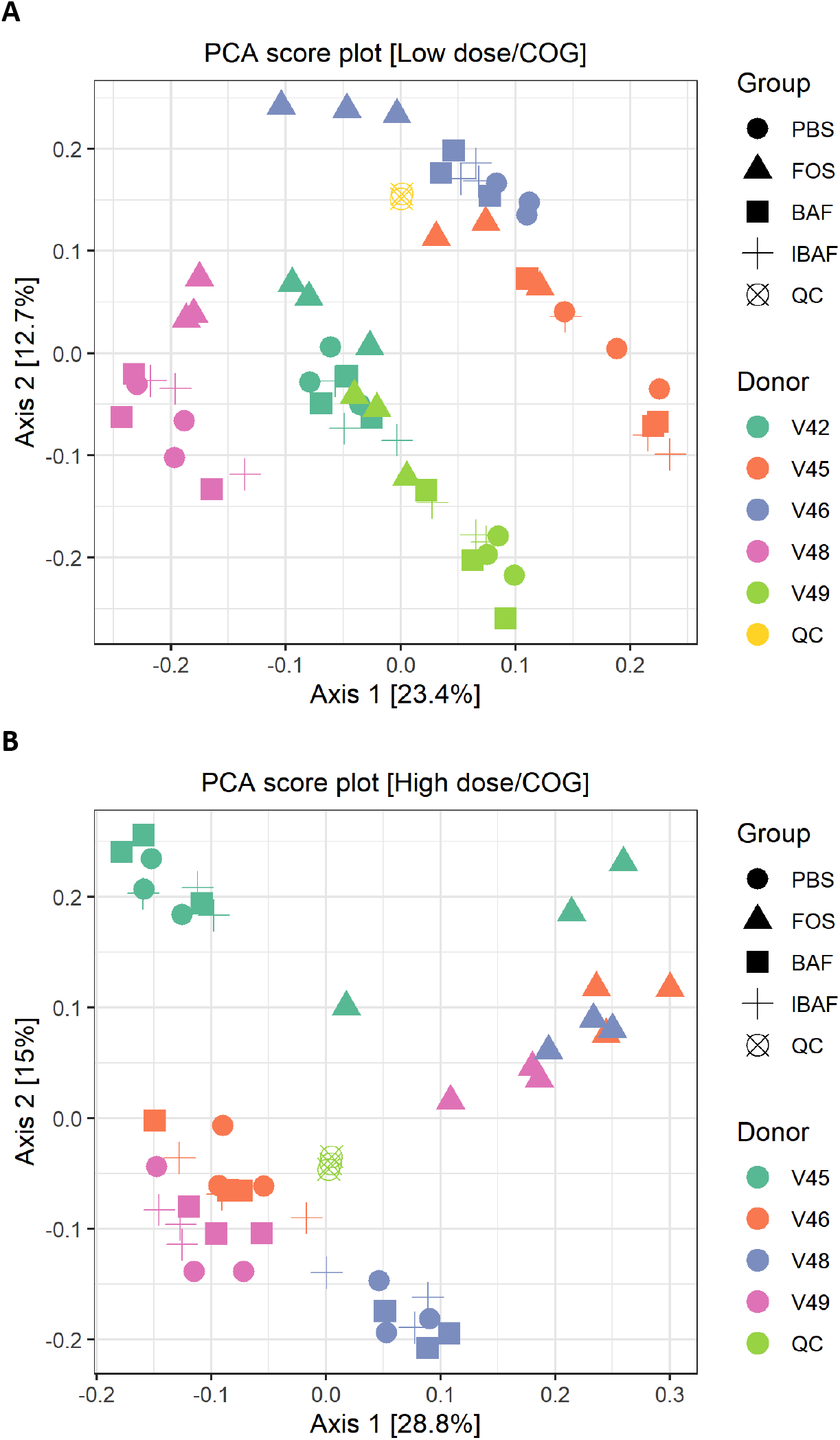
Principal component analyses of clusters of orthologous groups (COGs) demonstrate the unique pattern of an individual’s functional metaproteomic profile and absence of BAFASAL impact. Principal Component Analyses – COGs. Panels A and B show principal component analyses (PCA) using COGs per sample and for low and high doses, respectively. Colors indicate volunteer samples and shapes treatment as shown to legends on the right of each panel. Note: Only those COGs that have non-zero values in >50% of samples were used for analyses (Q50).

Supplemental Figures 6 to 8 illustrate the abundance distribution of major functional categories, including translation, amino acid metabolism, carbohydrate metabolism, lipid metabolism, and energy metabolism. There were no significant changes for any individual nor dosage when comparing BAF, IBAF, and PBS treated microbiomes for diverse COGs such as translation, ribosomal structure, and biogenesis; amino acid transport and metabolism; lipid transport and metabolism; nucleotide transport and metabolism; carbohydrate transport and metabolism; and energy production and conversion. In contrast, the FOS positive control had significant effects on multiple COG categories, including increased functional categories of translation and amino acid metabolism; decreased lipid transport and metabolism; increased nucleotide transport and metabolism; increased carbohydrate transport and metabolism, and increased energy production and conversion, all at low and high doses with inter-individual variation and responses.

### 3.5 Metaproteomic comparative taxonomic analyses

Additionally, and complementing our 16S analyses, we performed comparative taxonomic analyses at phylum and genus levels using quantified peptides in metaproteomics (Figure 8 – low dose and Supplemental Figure 9 – high dose). No phylum-level changes were observed among BAF, IBAF, and PBS treated microbiomes for an individual (Panel A). Several phyla abundances were significantly increased in FOS-treated versus vehicle PBS-treated microbiomes, including Actinobacteria, as expected and documented in the literature [27]. Similarly, no obvious genus-level changes were observed among BAF, IBAF, and PBS treated microbiomes for an individual. (Panel B). However, FOS treatment increased *Bifidobacterium* as expected and documented in the literature [27].

**Figure 8.**
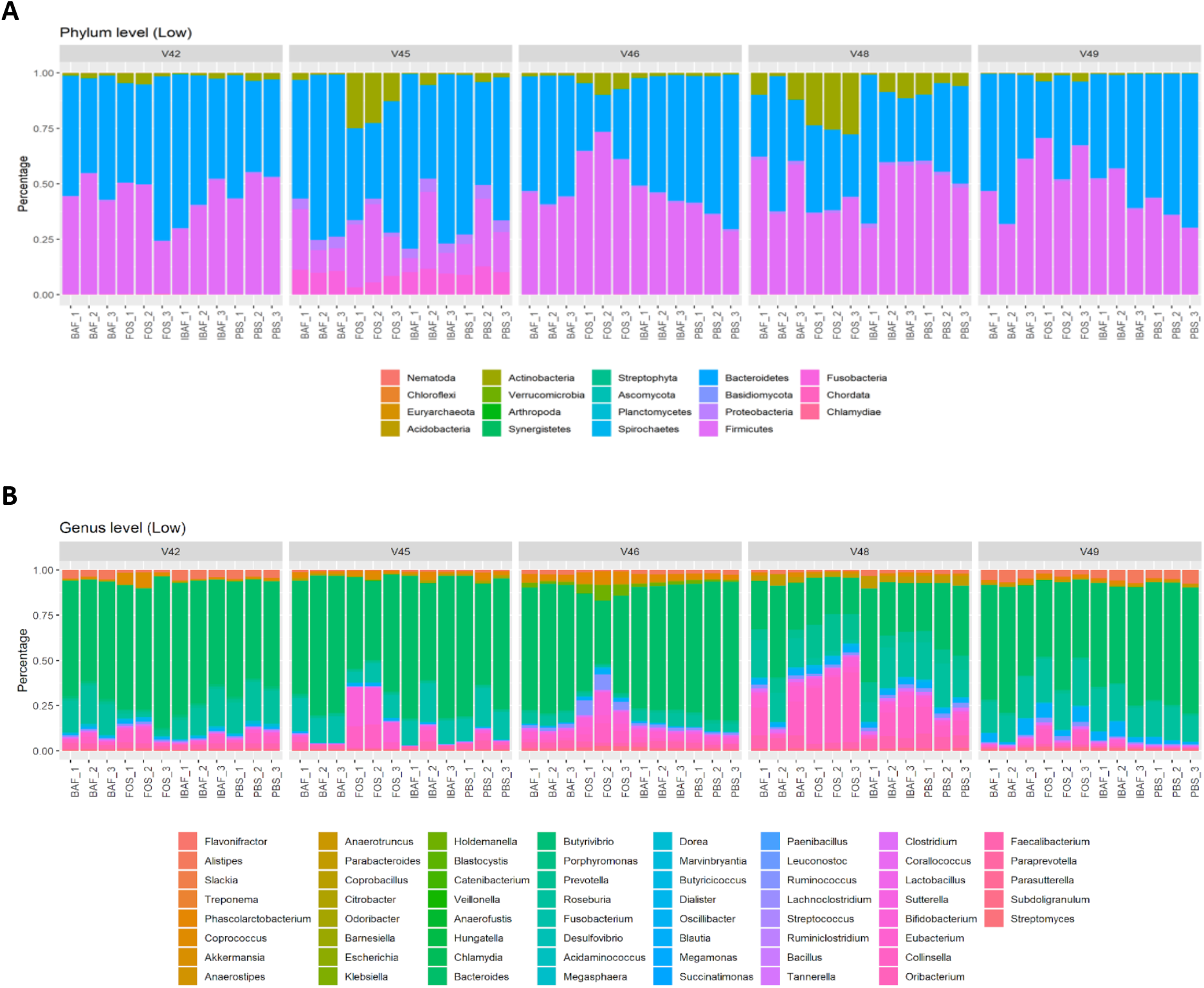
Taxonomic analyses at the phylum and genus levels demonstrate the unique composition of an individual’s microbiome via metaproteomic profiling and absence of BAFASAL impact. Stacked bar representation from label free quantification of protein groups assigned to phylum (Panel A – low dose and genus levels (Panel B – low dose) expressed as percentages. Individual microbiomes are grouped, and the phylum and genus are identified by colored bars as indicated in key below panels.

## 4. Discussion

This study aimed to examine the effects of the BAFASAL agricultural bacteriophage mixture on human gut microbiome function and composition using an *in vitro* RapidAIM platform. In particular, we were interested in whether this agricultural bacteriophage mixture, used as a feed additive in poultry farming to prevent and eliminate *Salmonella*, would display off-target effects on the human microbiome. It has been reported that BAFASAL significantly reduces *Salmonella* levels in poultry, reducing mortality and improving feed conversion rates [12]. These types of therapeutic and prophylactic agricultural bacteriophages are important tools to address the critical on-going need to maintain food safety, improve the health of animals in our food chains, and reduce human morbidity and mortality due to consumption of contaminated food. Bacteriophage development, and studies regarding their safe introduction into our food chain, have been prompted in light of growing antimicrobial resistance and the environmental pollution caused by decades-long wide-spread antibiotic use in the food industry [4].

Bacteriophages, as self-replicating and self-limiting agents, are generally regarded as safe. A bacteriophage preys on specific bacterial strains due to their specificity in recognizing particular receptors on target bacteria [7]. Additionally, virulent bacteriophages that exhibit only a lytic mechanism of replication, of which BAFASAL is composed, are desirable as therapeutics since once their target bacteria are infected and lysed, the bacteriophages themselves become self-limiting. To date, rodent models have been used extensively to examine bacteriophage safety as therapeutics in medicine and agriculture. Oral administration of different anti-*Salmonella* bacteriophage mixtures into mice did not result in gross clinical changes or mortality [12, 28]. Additionally, mice treated with a bacteriophage therapeutic against *E.coli* also did not exhibit any toxic effects [29].

Herein, we determined whether the bacteriophage mixture BAFASAL, proven effective against *Salmonella sp*. in poultry, would affect the human gut microbiome using our novel multi-omics RapidAIM approach against healthy adult microbiomes. 16S rRNA gene sequencing captured compositional changes and was complemented by proteomics analyses that, in addition to providing confirmatory information on taxonomic abundances, can accurately quantify expressed proteins and assess functional changes. It was important to use both approaches as our previous work demonstrated that bacterial function could change and shift in response to a stimulant without a change in the abundance of bacteria [20].

As a gross measure of BAF potential impact on gut microbiota, we showed that total biomass was unaffected compared to IBAF treated microbiomes (Figure 2). Nor did it differ from our PBS treated negative control. In contrast, FOS-treated microbiome biomasses were lower than all other groups, suggesting FOS treatment depresses selective or general gut bacterial growth. Microbial richness and diversity were unaffected by BAF treatments (Figure 3), while FOS treatment reduced microbial richness, likely due to specific microbes thriving when using this carbon source and suppressing the growth of competitors. Similarly, only FOS significantly affected beta-diversity metrics for each individual microbiome, while BAF treated microbiomes were indistinguishable from their corresponding heat-inactivated and PBS-treated microbiomes (Figures 4-5). Overall, metagenomics using 16S rRNA amplicon analyses did not reveal significant changes in the composition or abundances of the microbiomes obtained from any of the five healthy adults following 18hr culture with BAF at two concentrations in comparison to IBAF or PBS. These results differed from the FOS positive control, which resulted in the expected and documented increases in *Bifidobacterium* genera and reduced levels of several Enterobacteriaceae family members (Supplemental Figure 3)[27, 30].

Hierarchical clustering of microbial proteins did not reveal shifts in protein abundances among BAF, IBAF, and PBS treatments at either dose for any individual microbiome tested, while FOS treatment-induced changes in protein expression so that these samples clustered at both low and high doses. In fact, at high dose, FOS treatment protein abundance changes overcame intraindividual clustering (Supplemental Figure 4). Likewise, PCA analyses did not detect any shifts in protein differences in BAF, IBAF, and PBS treatments at either dose while low dose FOS treated microbiomes separated from their individual microbiome, and at high dose, those FOS-induced changes were increased to the level that would overcome interindividual differences (Figure 6). Collectively, these analyses did not reveal any significant shifts in protein abundance with BAF treatment. Results from FOS treatments established the utility of the metaproteomic branch of the RapidAIM assay to reliably and reproducibly detect expressed protein abundance changes and that dose-dependent effects can be measured and differentiated. Quantified protein groups were assigned to functional pathways to examine whether microbial function differs with BAF treatment (Figure 7). There were no differences detected under BAF treatment (Supplemental Figures 6-8), while FOS treated microbiomes show predicted changes from past metagenomic studies [27, 30]. Metaproteomic assessment of phylum and genus level abundances in the presence of BAF treatment were similar to IBAF, and PBS treated microbiomes. Overall, metaproteomic analyses did not reveal obvious protein abundance changes nor functional changes of the microbiomes obtained from any of the five healthy adults following 18hr culture with BAF at two concentrations and, in comparison to IBAF or PBS. Meanwhile, our control FOS showed the expected and documented changes such as increases in *Actinobacteria* and a decrease in *Proteobacteria* (Figure 8 and Supplemental Figure 9) [27, 30].

The gut microbiome composition of humans has a definite impact on health and disease. The gut is an important milieu where the host, pathogens, and foods interact. Shifts in our microbiome due to diet, treatments, therapies, antibiotics can be correlated with diseases. As such, food industry leaders and government regulators are closely studying the impact of food processing, additives, etc., on the microbiome. Antibiotics, especially broad-spectrum antibiotics, have profound effects on the gut microbiota. We have recapitulated and documented these effects previously using the RapidAIM platform [20]. Bacteriophages are very abundant in human guts as stable members of our niche microbiota, with estimates of 10^12^ bacteriophages per gram found in our feces. Many are lysogenic phages, and replication only occurs in their bacterial hosts. Nonetheless, it is vital to understand whether bacteriophages introduced and used as therapies in, for instance, agriculture, could make their way into the human intestinal system and have significant off-target effects. Agricultural workers may be uniquely at risk for any effects of these new therapies and prophylactics as they may be exposed to agricultural residues with residual bacteriophage from the feed.

The RapidAIM platform as utilized in this study provided an opportunity to study the effects of the bacteriophage mixture BAFASAL on human gut microbiota and on an individual basis. To our knowledge, this is the first study of this kind to measure the response of the human gut microbiome to bacteriophage-based product. That we measure responses on an individual basis is a strength of this *in vitro* assay as studies show our microbiomes are as unique as their hosts and can respond differentially to stimuli. Indeed, we have previously how individual microbiomes vary in their response to drug treatments [20]. We examined five individual microbiomes’ response to BAFASAL in this pioneering study; however, the RapidAIM platform is scalable to large numbers of individuals, can be modified for pooled effects, and can simultaneously screen large numbers of compounds. In this study, we demonstrated that BAFASAL did not affect microbiome composition nor microbiome function from the five individual participants and suggested that results from such an assay can be added to therapeutic and prophylactic bacteriophage safety profiling.

## Conclusions

The goal of our study was to utilize RapidAIM, a novel *in vitro* assay of the gut microbiome, to assess the effect of BAFASAL bacteriophage preparation on our gut microbiota. In this multi-omics study, including 16S rRNA gene sequencing and metaproteomics, we show that BAFASAL does not affect healthy human adult microbiomes’ composition and function. This study also highlights the value of the RapidAIM assay as a tool/platform to rapidly measure whether, or not novel antibacterial solutions, including bacteriophage-based products, affect our niche gut microbiota – ecology.

## Acknowledgments

The authors wish to thank the volunteers who participated in this study. This research received no external funding.

## Author Contributions

All authors participated in the design of this study. JM coordinated the volunteers, and JM, XZ, and KW collected the samples. JM, XZ, JB, KW and ZN developed the workflow and performed the experiments. XZ and JB analyzed and visualized the data. JM, DF, JB, XZ, and AS wrote the manuscript. All authors participated in data interpretation, edited and approved the final manuscript.

## Conflicts of Interest

DF and AS have co-founded Biotagenics and MedBiome, clinical microbiomics companies. JD is a founder of the company Proteon Pharmaceuticals S.A., which developed the BAFASAL preparation. All other authors declare no potential conflicts of interest.

## Supplemental Figures

**Supplemental Figure 1: RapidAIM assay and sequencing pipeline generate reproducible results.** Principal coordinates analysis (PCoA) using the Bray-Curtis dissimilarity revealed that the individual stools clustered tightly together (A) with essentially no separation between the two sequencing chips. In addition, the PBS replicates from the high/low dose conditions were also highly similar (B), with the samples segregating primarily by the donor. Our commercial standard was also highly similar between sequencing runs and could distinguish between the related *Escherichia coli* and *Salmonella enterica* (C). There were some reads aligning to Enterobacteriaceae genus *Enterobacter*, which appeared to be primarily derived from *Escherichia* reads.

**Supplemental Figure 2: Principal coordinate analysis of RapidAIM cultures from each donor reveals that BAFASAL treatment has minimal impact on microbial composition.** PBS/BAF/IBAF RapidAIM assays from each donor were analyzed separately using principal coordinate analysis of the Bray-Curtis dissimilarity and plotted as separate panels. There were no apparent differences between the PBS and BAFASAL/inactivated BAFASAL treatments even after removing the FOS samples.

**Supplemental Figure 3: BAFASAL has no apparent impact on the abundances of Enterobacteriaceae genera.** The log10 Relative Abundances for the five most abundant Enterobacteriaceae genera are plotted for each treatment (high/low merged). A pseudo count of 1 read was added to samples with zero abundances to allow for log_10_ plotting. There was no difference between the PBS/BAF/IBAF treatments, with several Enterobacteriaceae genera showing reductions under FOS treatment.

**Supplemental Figure 4: Sample clustering at the protein level does not support an effect for BAFASAL on human microbiome protein expression.** Panels A and B show sample clustering at the protein level per sample and for low and high doses, respectively. Colors indicate quality control (QC) or volunteer and treatment, as shown to legends on the right of the cluster figure. Colors within the cluster diagram indicate protein expression level, and black lines on the left of the figure indicate protein groups. Note: Only those protein groups with non-zero values in >50% of samples were used for analyses (Q50). Volunteer’s 42 high dose treatment results were removed from further analyses as we did not have qualifying replicates for its FOS group.

**Supplemental Figure 5: Sample clustering at the functional level does not support an effect for BAFASAL on the human microbiome.** Panels A and B show clustering of orthologous groups at the protein level per sample and for low and high doses, respectively. Colors indicate quality control (QC) or volunteer and treatment, as shown to legends on the right of the cluster figure. Colors within the cluster diagram indicate protein expression level, and black lines on the left of the figure indicate protein groups. Note: Only those COGs with non-zero values in >50% of samples were used for analyses (Q50). Volunteer’s 45 high dose FOS 3 was an outlier (V45_FOS_3).

**Supplemental Figure 6: Abundance distribution of major functional categories via clusters of orthologous groups (COGs) analyses for translation, ribosomal structure and biogenesis, and amino acid transport and metabolism.** Box and Whisker plots from label-free quantification of protein groups assigned to functional COGS translation, ribosomal structure and biogenesis (Panels A – low dose and C – high dose), and amino acid transport and metabolism (Panels B – low dose and D – high dose). Individual microbiomes are grouped, and the treatment is indicated by color according to the key to the right of the panels.

**Supplemental Figure 7: Abundance distribution of major functional categories via clusters of orthologous groups (COGs) analyses for lipid transport and metabolism, and nucleotide transport and metabolism.** Box and Whisker plots from label-free quantification of protein groups assigned to functional COGS of lipid transport and metabolism (Panels A – low dose and C – high dose) and nucleotide transport and metabolism (Panels B – low dose and D – high dose). Individual microbiomes are grouped, and the treatment is indicated by color according to the key to the right of the panels.

**Supplemental Figure 8: Abundance distribution of major functional categories via clusters of orthologous groups (COGs) analyses for carbohydrate transport and metabolism, and energy production and conversion:** Box and Whisker plots from label-free quantification of protein groups assigned to functional COGS of carbohydrate transport and metabolism (Panels A – low dose and C – high dose) and energy production and conversion (Panels B – low dose and D – high dose). Individual microbiomes are grouped, and the treatment is indicated by color according to the key to the right of the panels.

**Supplemental Figure 9. Taxonomic analyses at the phylum and genus levels demonstrate the unique composition of an individual’s microbiome via metaproteomic profiling and absence of BAFASAL impact.** Stacked bar representation from label free quantification of protein groups assigned to phylum (Panel A – high dose) and genus levels (Panel B – high dose) expressed as percentages. Individual microbiomes are grouped, and the phylum and genus are identified by colored bars as indicated in key below panels.

