## Supplemental Data Figures for "Examining the effects of an anti-Salmonella bacteriophage preparation, BAFASAL, on *ex vivo* human gut microbiome composition and function using a multi-omics approach"

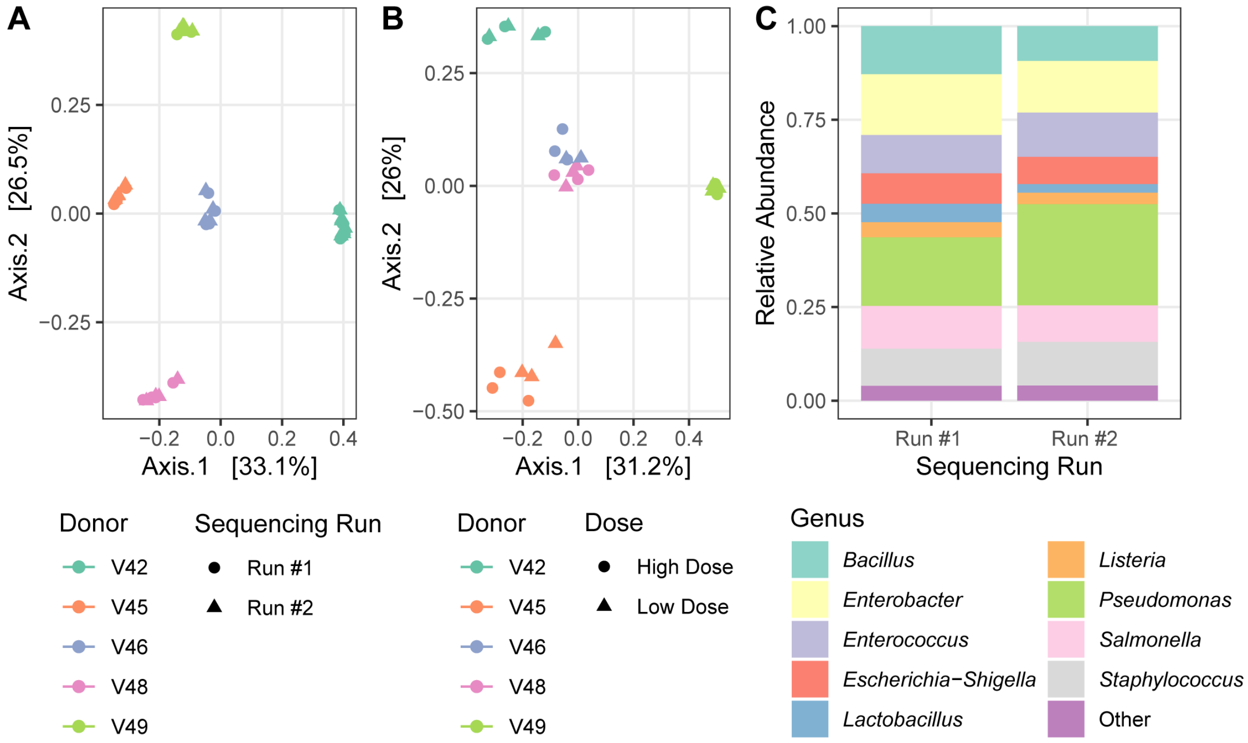


**Supplemental Figure 1: RapidAIM assay and sequencing pipeline generate reproducible results.** Principal coordinate analysis (PCoA) using the Bray-Curtis dissimilarity revealed that the individual stools clustered tightly together (A) with essentially no separation between the two sequencing chips. In addition, the PBS replicates from the high/low dose conditions were also highly similar (B), with the samples segregating primarily by donor. Our commercial standard was also highly similar between sequencing runs and could distinguish between the related *Eschericha coli* and *Salmonella enterica* (C). There were some reads aligning to Enterobacteriaceae genus *Enterobacter* which appeared to be primarily derived from *Escherichia* reads.

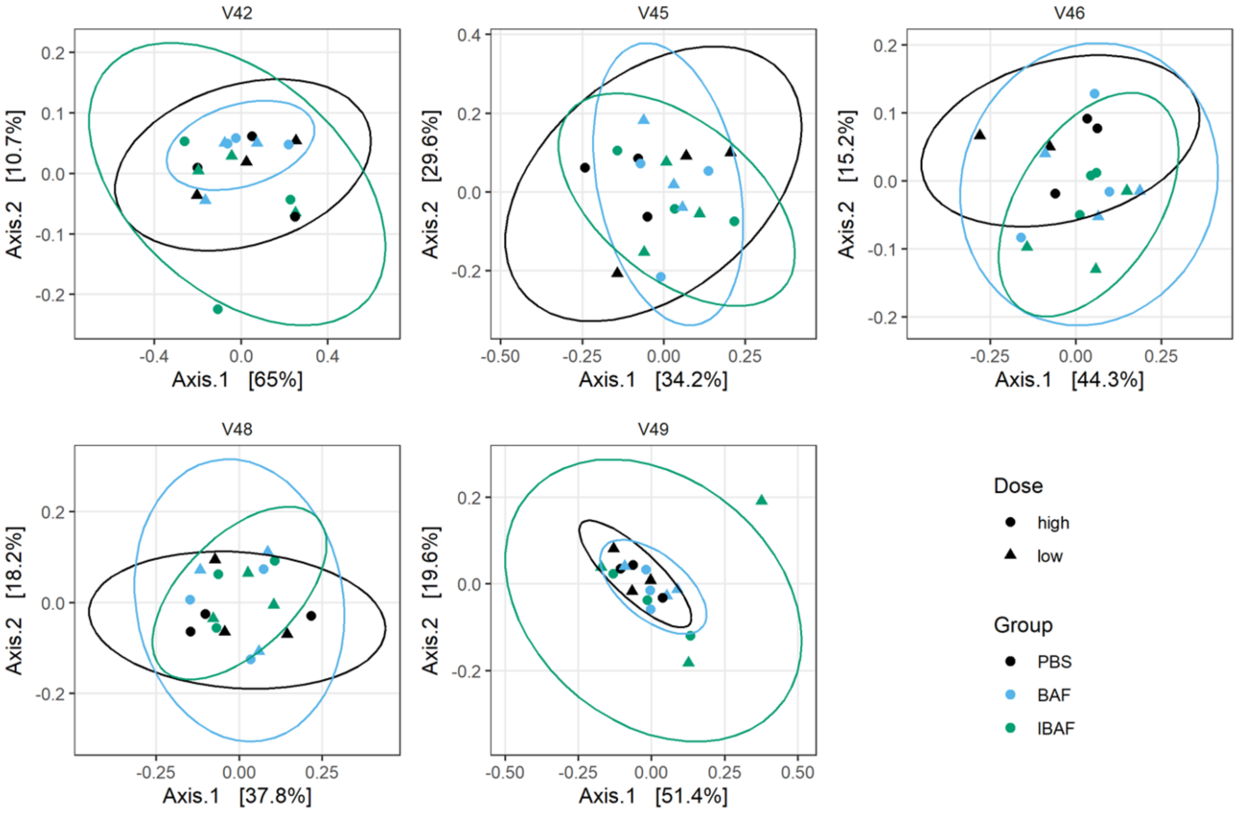


**Supplemental Figure 2: Principal coordinate analysis of RapidAIM cultures from each donor reveals that BAFASAL treatment has minimal impact on microbial composition.** PBS/BAF/IBAF RapidAIM assays from each donor were analyzed separately using principal coordinate analysis of the Bray-Curtis dissimilarity and plotted as separate panels. There were no apparent differences between the PBS and BAFASAL/inactivated BAFASAL treatments even after removing the FOS samples.

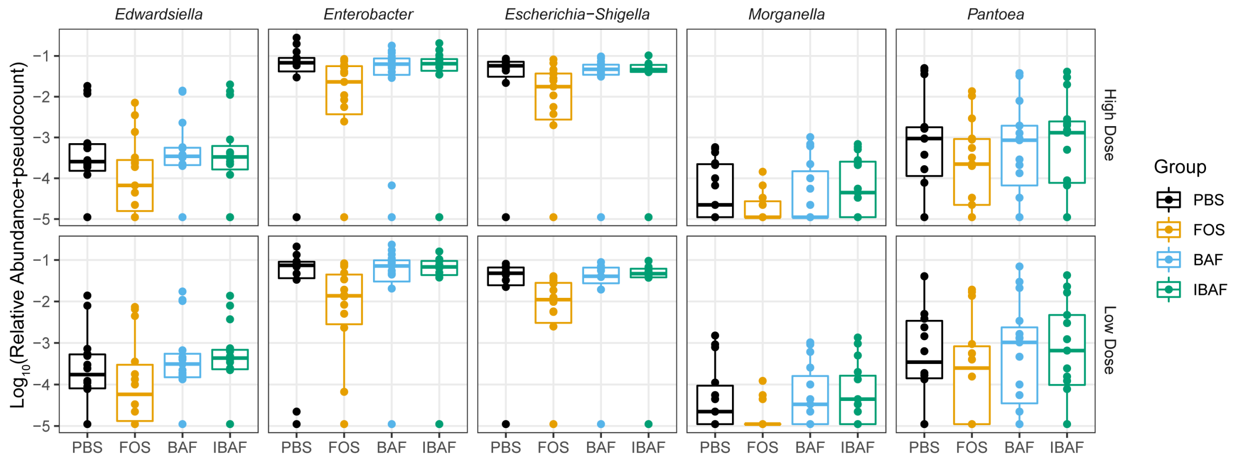


**Supplemental Figure 3: BAFASAL has no apparent impact on the abundances of Enterobacteriaceae genera.** The log10 Relative Abundances for the 5 most abundant Enterobacteriaceae genera are plotted for each treatment (high/low merged). A pseudocount of 1 read was added to samples with zero abundances to allow for log10 plotting. There was no difference in between the PBS/BAF/IBAF treatments with several Enterobacteriaceae genera showing reductions under FOS treatment.

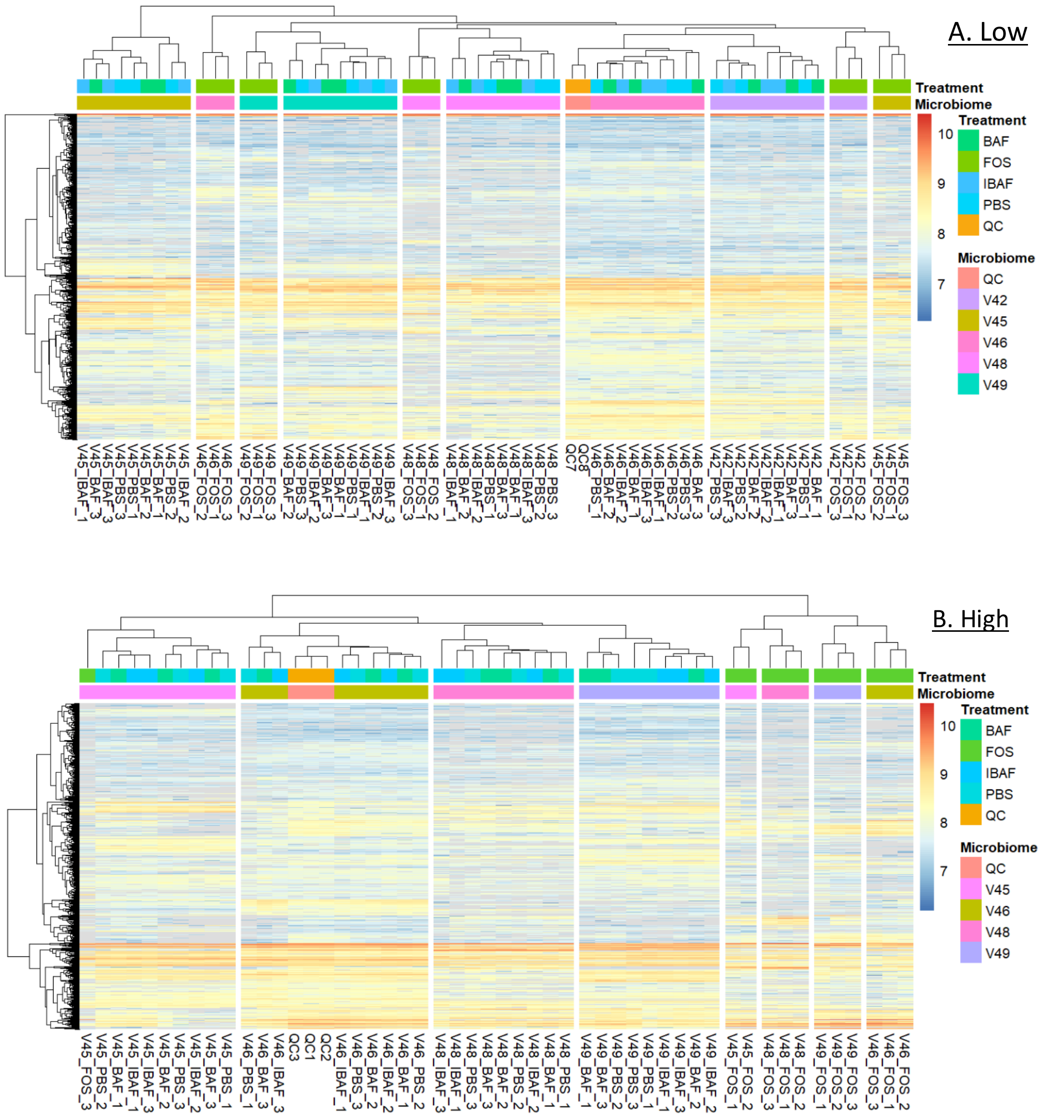


**Supplemental Figure 4: Sample clustering at the protein level does not support an effect for BAFASAL on human microbiome protein expression**. Panels A and B show sample clustering at the protein level per sample and for low and high doses, respectively. Colors indicate quality control (QC) or volunteer and treatment as shown to legends on the right of cluster figure. Colors within cluster diagram indicate protein expression level and black lines on left of figure indicate protein groups. Note: Only those protein groups that have non-zero values in >50% of samples were used for analyses (Q50). Volunteer's 42 high dose treatment results were removed from further analyses as we did not have qualifying replicates for it's FOS group.

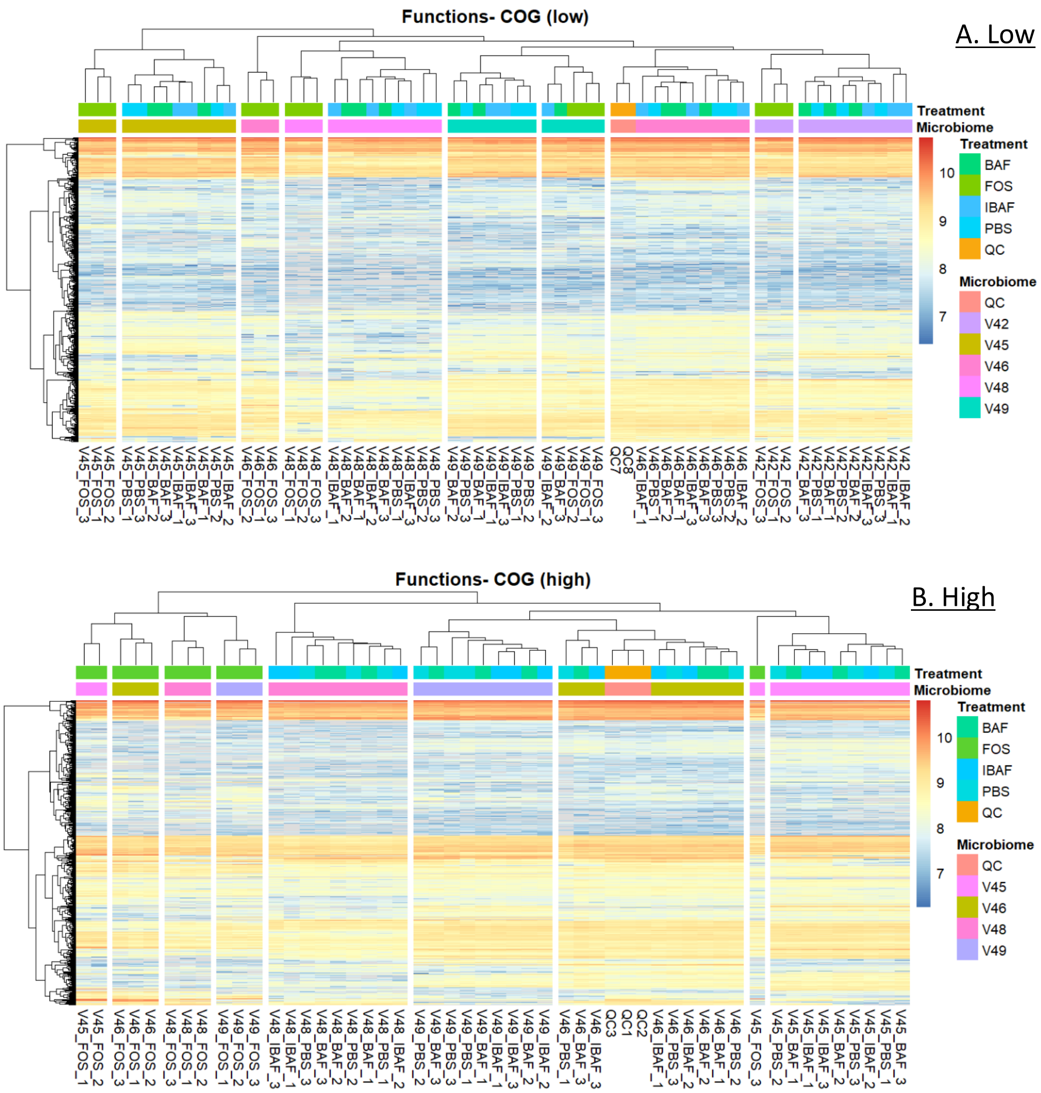


**Supplemental Figure 5**: **Sample clustering at the functional level does not support an effect for BAFASAL on the human microbiome**. Panels A and B show clustering of orthologous groups at the protein level per sample and for low and high doses, respectively. Colors indicate quality control (QC) or volunteer and treatment as shown to legends on the right of cluster figure. Colors within cluster diagram indicate protein expression level and black lines on left of figure indicate protein groups. Note: Only those COGs that have non-zero values in >50% of samples were used for analyses (Q50). Volunteer's 45 high dose FOS 3 was an outlier (V45_FOS_3).

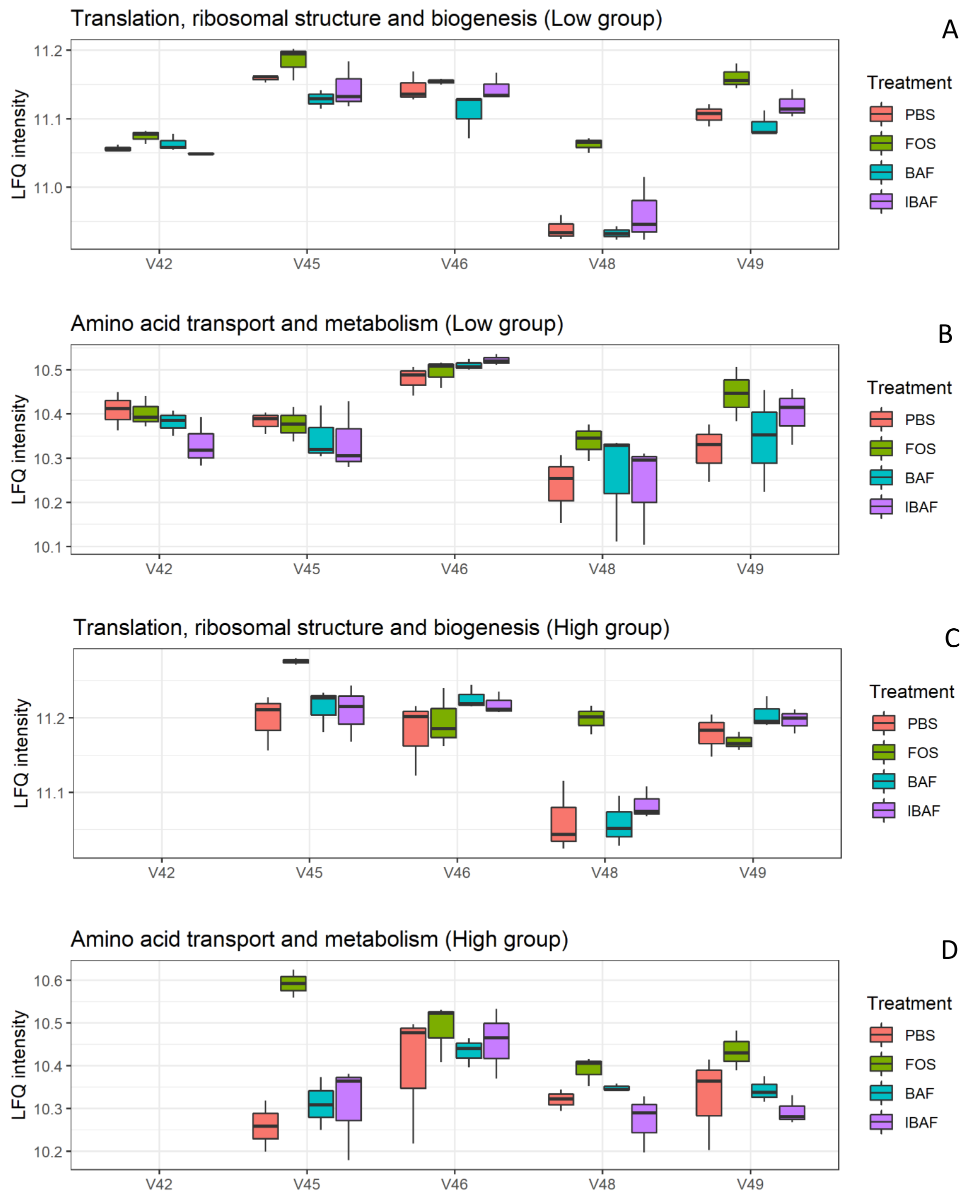


**Supplemental Figure 6: Abundance distribution of major functional categories via clusters of orthologous groups (COGs) analyses for translation, ribosomal structure and biogenesis, and amino acid transport and metabolism**. Box and Whisker plots from label free quantification of protein groups assigned to functional COGS translation, ribosomal structure and biogenesis (Panels A – low dose and C – high dose) and amino acid transport and metabolism (Panels B – low dose and D – high dose). Individual microbiomes are grouped, and the treatment is indicated by color according to key to right of panels.

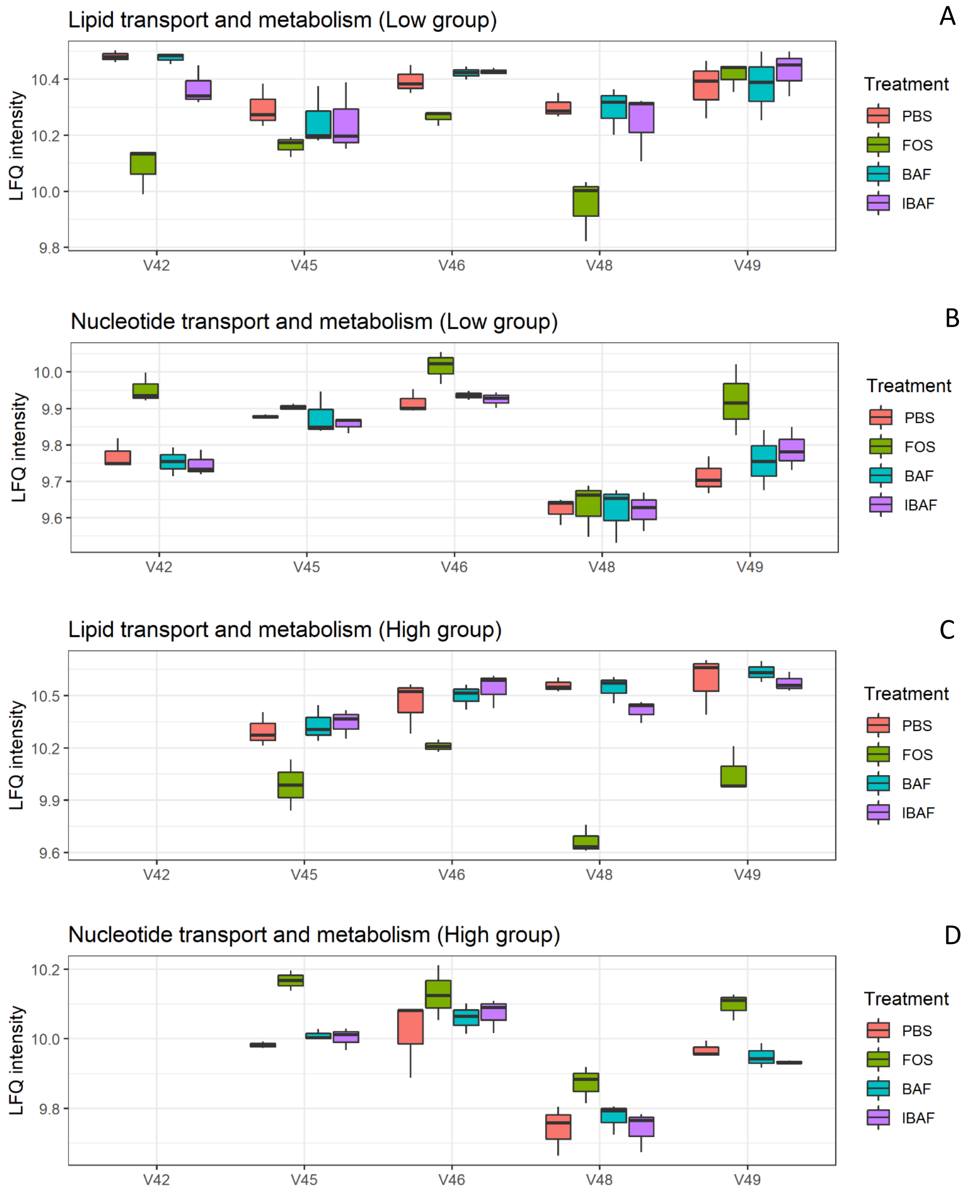


**Supplemental Figure 7: Abundance distribution of major functional categories via clusters of orthologous groups (COGs) analyses for lipid transport and metabolism, and nucleotide transport and metabolism**. Box and Whisker plots from label free quantification of protein groups assigned to functional COGS of lipid transport and metabolism (Panels A – low dose and C – high dose) and nucleotide transport and metabolism (Panels B – low dose and D – high dose). Individual microbiomes are grouped, and the treatment is indicated by color according to key to right of panels.

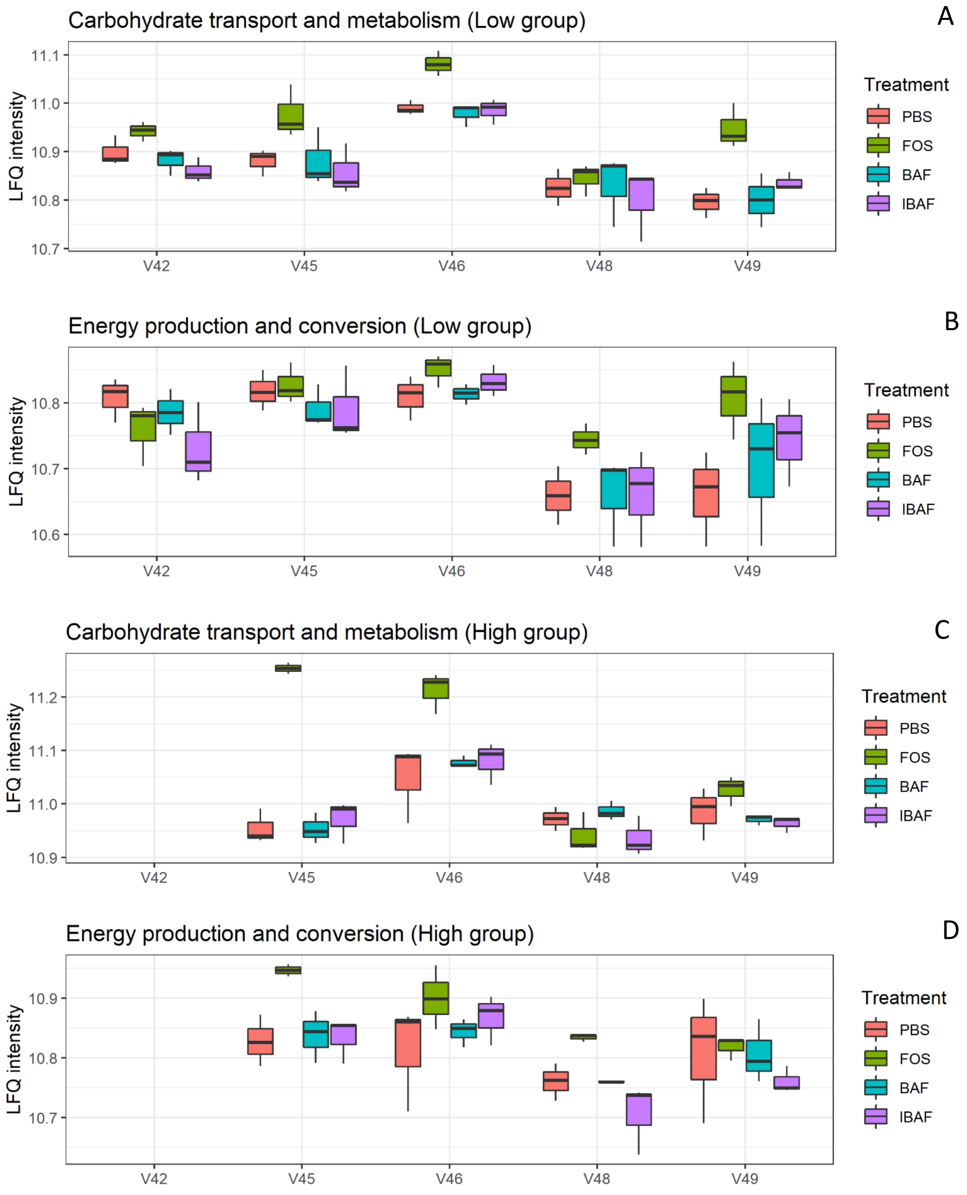


**Supplemental Figure 8: Abundance distribution of major functional categories via clusters of orthologous groups (COGs) analyses for carbohydrate transport and metabolism, and energy production and conversion:** Box and Whisker plots from label free quantification of protein groups assigned to functional COGS of carbohydrate transport and metabolism (Panels A – low dose and C – high dose) and energy production and conversion (Panels B – low dose and D – high dose). Individual microbiomes are grouped, and the treatment is indicated by color according to key to right of panels.

A

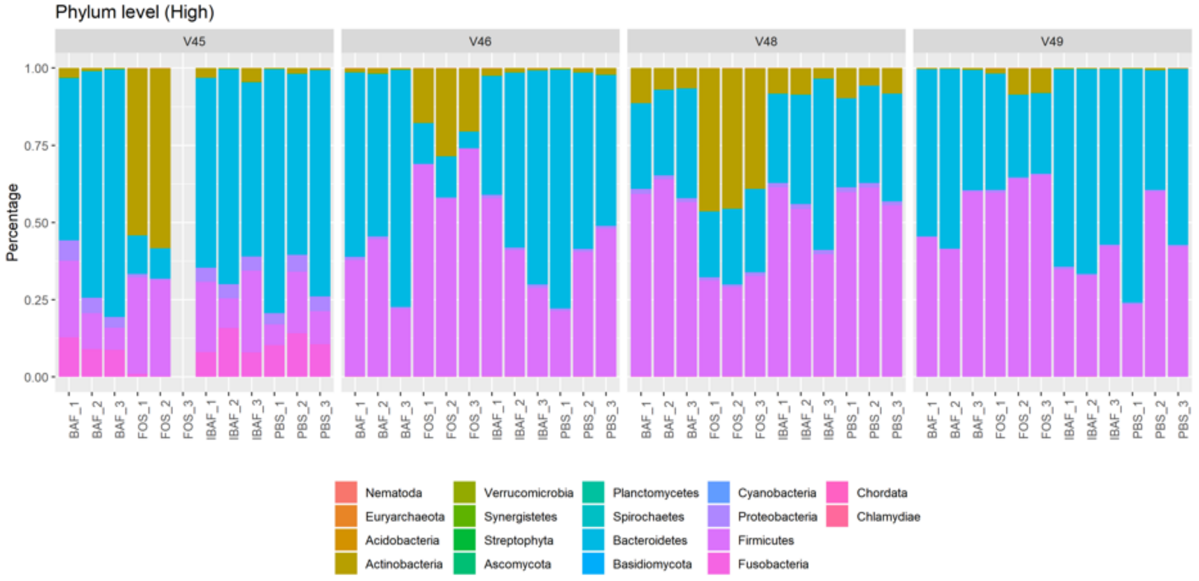


B

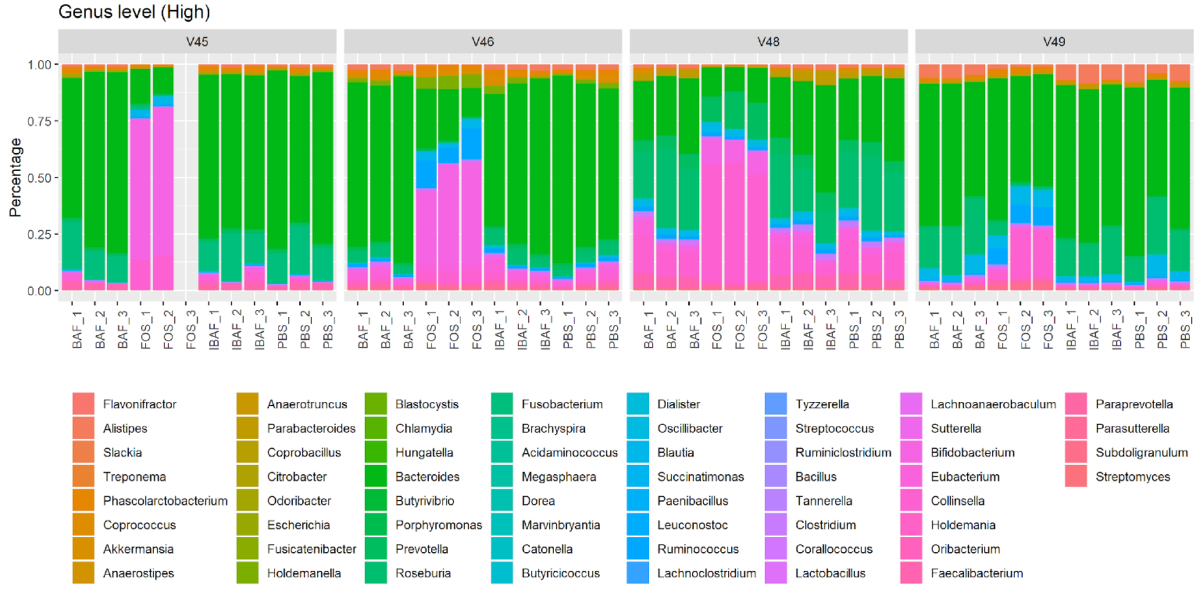


**Supplemental Figure 9. Taxonomic analyses at the phylum and genus levels demonstrate the unique composition of an individual's microbiome via metaproteomic profiling and absence of BAFASAL impact.** Stacked bar representation from label free quantification of protein groups assigned to phylum (Panel A – high dose) and genus levels (Panel B – high dose) expressed as percentages. Individual microbiomes are grouped, and the phylum and genus are identified by colored bars as indicated in key below panels.
